# Structure-informed evolutionary analysis of the meiotic recombination machinery

**DOI:** 10.1101/2025.11.07.687317

**Authors:** Meret Arter, Jeffrey Vedanayagam, Min Lu, Momar Diop, Kaixian Liu, Eric C. Lai, Scott Keeney

**Author notes:** Department of Reproductive Endocrinology, University Hospital Zurich, University of Zurich, 8952 Schlieren, Switzerland (Meret Arter). Department of Neuroscience, Developmental and Regenerative Biology, University of Texas at San Antonio, San Antonio, TX 78249. **Corresponding Author: Scott Keeney Email:**. **Author Contributions:** M.A., J.V., and S.K. designed research; M.A., J.V., M.L., M.D., and K.L. performed research; M.A., J.V., M.L., M.D., K.L, E.C.L. and S.K. analyzed data; and M.A., J.V. and S.K. wrote the paper.

## Abstract

Despite being essential for fertility, many proteins involved in meiotic homologous recombination have diverged rapidly. The evolutionary forces driving this divergence remain mostly unknown, in part because of challenges in accounting for the interplay of sequence changes with constraints imposed by proteins’ structures and physiological roles. Here, we explore strategies to more sensitively detect signatures of positive or relaxed selection by integrating evolutionary analyses with structural and functional information, using meiotic recombination proteins in four taxa—primates, rodents, birds and budding yeasts. By mapping selection rate estimates onto predicted protein structures, we characterized protein regions likely to have experienced positive selection. We further identified subtle sequence variation within protein domains that are well conserved generally because of structural constraints. To detect sequence variation masked by these constraints, we analyzed selection at structurally matched residues, comparing homologs across different lineages as well as between meiosis-specific and generalist paralogs. These approaches identified lineage- and paralog-restricted enrichment of non-synonymous substitutions that may indicate loss of functional constraints and/or adaptive innovation. Finally, we used cross-species complementation experiments in *Saccharomyces cerevisiae* to show that sequence variation in the pro-crossover factor MSH4 modulates recombination proficiency. We suggest that evolutionary plasticity per se is a key conserved characteristic of the meiotic recombination machinery. More generally, our approach provides a mechanistic framework to analyze protein evolution.

**Significance Statement:** Meiotic recombination—the shuffling of genetic material during reproductive cell formation—is essential for fertility across all sexually reproducing organisms. Paradoxically, the proteins mediating recombination evolve rapidly despite their critical function. We used an integrative approach combining evolutionary analysis with protein structure prediction to understand how and why these proteins change. By mapping sequence variation onto protein structures and comparing meiosis-specific proteins to their generalist counterparts, we identified regions likely experiencing adaptive evolution or relaxed functional constraints. Experimental tests confirmed that sequence differences affect function. Our findings suggest that evolutionary plasticity itself may be a conserved feature of the recombination machinery. This structure-informed framework offers a generalizable strategy for detecting functionally relevant evolutionary changes in other protein systems.

## Introduction

To keep genome content stable across generations, sexually reproducing eukaryotes generate gametes that contain only half the parent’s chromosome number. Most organisms generate gametes through meiosis, a specialized cell division in which maternal and paternal homologous chromosomes usually use homologous recombination to pair, exchange genetic information, and then subsequently segregate (1, 2).

Meiotic recombination is initiated by DNA double-strand breaks (DSBs) generated by topoisomerase-like SPO11 and its partners (3) (**Fig. 1*A***). DSBs are resected by endo- and exonucleases (4). The resulting long single-stranded 3’ DNA overhangs are bound by ssDNA binding proteins and RAD51-like recombinases that mediate invasion of the homologous chromosome to form branched recombination intermediates including D-loops, single-end-invasions, and double Holliday junctions (5). Two main pathways result in either noncrossover or crossover outcomes. Noncrossovers form mostly through synthesis-dependent strand annealing, involving the unwinding of early recombination intermediates by DNA helicases (6, 7). Crossovers are mostly formed through the asymmetric resolution of double Holliday junctions by nucleases (8). Only crossovers contribute to physical linkages between homologous chromosomes that are needed for segregation, but all recombination events contribute to chromosome pairing in many species, and all DSBs need to be faithfully repaired to maintain genomic integrity in all species. Meiotic recombination is an ancient process that is shared across eukaryotes and has been studied in many organisms. Meiotic recombination errors are generally associated with missegregation of chromosomes, which can lead to aneuploid gametes and infertility (9). While the process overall is widely conserved, functional and genomic studies repeatedly revealed substantial variability within the protein machinery that performs each step (10). Given that this machinery is important for the generation of viable gametes in the organisms studied, it is reasonable to assume that it was likewise important in their ancestors. Yet many of these proteins are poorly conserved, showing recurrent loss of individual proteins or protein families, emergence of lineage-specific proteins, changes in the meiotic function of conserved proteins, and/or evolutionarily rapid changes in amino acid sequences (10).

**Figure 1.**
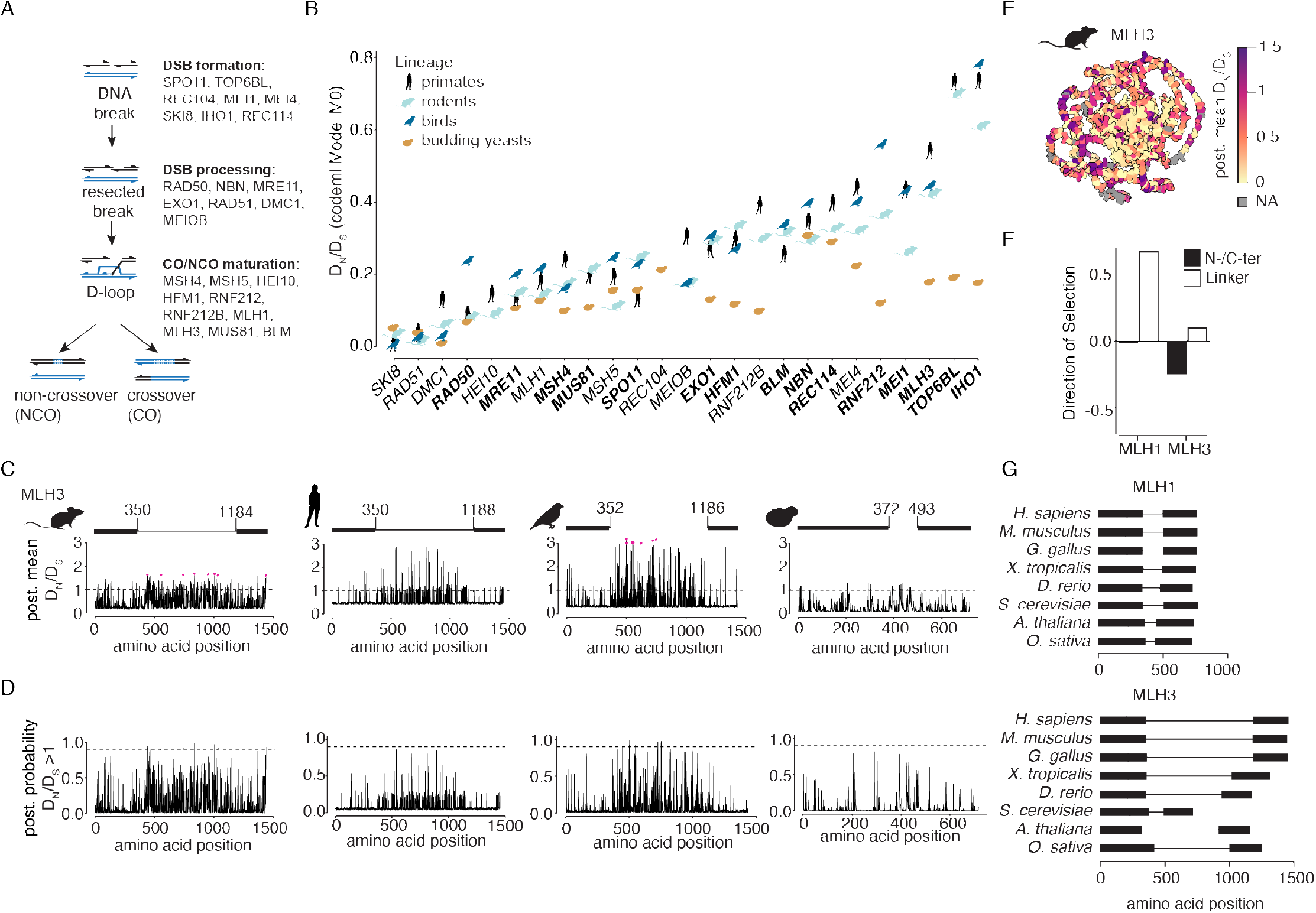
Structure-informed evolutionary analyses of the meiotic recombination machinery. (*A*) Conserved DNA events and proteins required for meiotic recombination (*B*) Average D_N_/D_S_ ratio for each gene. The genes are ordered according to the average across the four lineages. Genes in bold showed significant signs of positive selection in at least one lineage (LRT for codeml site models M7 vs. M8). (*C,D*) BEB estimation of post. mean D_N_/D_S_ values in panel *C* and post. probability for positive selection in panel *D* for each residue in MLH3 in the four lineages. Residues with a post. probability > 0.9 for positive selection are highlighted in pink in panel *C*. Dashed lines indicate D_N_/D_S_ = 1 in panel *C* and post. probability = 0.9 in panel *D*. The schematic above each graph in panel *C* shows positions of the N-and C-terminal folded domains (thick lines, from Interpro) and the central, unstructured linker (thin lines). (*E*) Post. mean D_N_/D_S_ mapped onto the AlphaFold model of mouse MLH3. The color scheme for mean D_N_/D_S_ values is capped at 1.5. (*F*) Direction of selection analysis in house mouse population data for the folded domains and unstructured linker regions of MLH1 and MLH3. (*G*) Greater variation between organisms in length of the unstructured linker in MLH3 than in MLH1.

It remains unclear how and why the meiotic recombination machinery is so evolutionarily plastic while continuing to maintain functionality over millions of years. An important step in answering this question is to understand at what rate and strength natural selection (purifying, neutral, or positive) is driving the evolution of these proteins. Using phylogenetic approaches, previous studies found signs of positive selection in several meiotic recombination proteins in different lineages such as flies, mammals or plants (11-15). However, most of these studies examined only a subset of the meiotic recombination machinery and they did not address which protein domains are most rapidly evolving.

Current strategies for detecting positive selection tend to work best for residues where recurrent positive selection leads to multiple amino acid substitutions; these strategies are often underpowered to detect other signatures of adaptive selection because of key limitations (16). First, selective pressures will generally not be all-purifying or all-positive: even amino acid residues under positive selection are also likely to have constraints on how freely they can change while maintaining the protein’s function. Second, single codon analysis of selection has limited statistical power, but increasing the power by averaging over multiple residues or entire proteins comes at the cost of potentially mixing residues with different combinations of selection (positive, purifying, neutral). Third, treating proteins as strings of amino acids without regard to three-dimensional structure or biochemical function can obscure evolutionary signatures. Finally, experimental studies often sidestep evolutionary variability entirely by focusing on the most conserved proteins or protein domains.

To address these issues, we explored strategies for increasing the sensitivity for detecting signatures of positive or relaxed selection by combining systematic evolutionary genetic analysis with structural and other biochemical information about the target proteins’ functions. We focused on core components of the meiotic recombination machinery required for DSB formation, end resection, strand invasion, and processing of recombination intermediates into crossover and noncrossover products. Our results yield evidence for otherwise cryptic positive selection in multiple phylogenetic lineages and suggest hypotheses about which protein domains and molecular functions have been the focus of evolutionary change.

## Results

### Overview of approach

The codeml package of Phylogenetic Analysis using Maximum Likelihood (PAML) (17) estimates synonymous (D_S_) and non-synonymous (D_N_) substitution rates. Under the simplifying assumption that synonymous substitutions are unseen by selection, a relative enrichment of non-synonymous substitutions (higher D_N_/D_S_ ratio) is a signature of reduced purifying selection often interpreted as positive selection. Codeml is a codon-based analysis that can be combined with protein structural models to generate further insights into which protein domains show signs of positive selection (18, 19). The advent of highquality structure prediction algorithms (20, 21) greatly expands the ability to integrate PAML with structural information. A recent study used this approach to infer functionally relevant sites under positive selection in centromere proteins (22).

Here we analyze meiotic recombination proteins in four eukaryotic lineages: primates, rodents, birds, and *Saccharomyces* budding yeasts (***SI Appendix* S1*A*-*D*; Dataset S1**). We integrated empirical or AlphaFold-generated structural models with PAML to determine how signatures of selection are shaped by structural constraints (folded domains vs. intrinsically disordered regions; buried/structural residues vs. protein-protein interfaces vs. solvent-exposed surfaces). We also sought to further disentangle positive selection signatures from the obscuring effects of purifying selection attributable to biochemical and structural constraints by comparing homologous proteins between different lineages and comparing paralogous protein pairs within lineages, where one paralog is (mostly) meiosis specific.

We selected 25 proteins that are directly involved in meiotic recombination (**Fig. 1*A***) and mined Ensembl, NCBI Refseq, and UniProt databases and used blastn searches of published genome assemblies to identify high-confidence gene sequences. In total, we compiled 1156 sequences encoding 25 proteins from 58 species in 4 lineages (***SI Appendix* Table S1 and Fig. S1*A*-*D***). For each lineage, we used MUSCLE (23) to generate amino acid-based multiple sequence alignments that were then manually curated (see ***SI Appendix Materials and Methods***).

### Many meiotic recombination proteins show signatures of positive selection and/or relaxed selection

We first ran codeml Model 0, implemented in the PAML package (17), to estimate an average D_N_/D_S_ ratio for the entire coding sequence of each protein in each lineage dataset using species trees (**Fig. 1*B* and *SI Appendix* Fig. S1*E***,***F***). Model 0 is a baseline model that estimates one D_N_/D_S_ ratio averaged across all sites and all species in the phylogeny. Genes with the highest overall D_N_/D_S_ were *IHO1, TOP6BL, MLH3*, and *RNF212*, while *SKI8, RAD51*, and *DMC1* showed the lowest ratio, reflecting high sequence conservation. PAML and similar algorithms have been widely used to detect signatures of positive selection (24).

To examine selection within different domains, we next ran codeml site models, which allow the D_N_/D_S_ ratio to vary across the gene and therefore allow detection of regions that are under positive selection in a gene that is overall under purifying selection. Site models treat the D_N_/D_S_ ratio for any codon in the gene as a random variable from a statistical distribution and different models use different underlying statistical distributions for D_N_/D_S_. Two pairs of nested models are most frequently used to test for sites under positive selection: models M1 (also called nearly neutral model) and M2 (positive selection), and models M7 (beta) and M8 (beta&ω). Only M2 and M8 allow for sites under positive selection. Likelihood ratio tests (LRTs) can evaluate which model better explains observed sequence variation.

PAML statistics of all LRTs are summarized in **Dataset S2**. No proteins scored as significant in yeast. However, synonymous substitutions were approaching saturation for many of the pairwise sequence comparisons (***SI Appendix*, Fig. S1*G***), which likely further limited statistical power. Therefore, failing to detect positive selection should not be viewed as evidence that it has not occurred. In contrast, the rodent, bird, and primate datasets had 15, 7, and 6 proteins under positive selection, respectively. The following proteins had codons with significant evidence of positive selection in at least one lineage (testing M1 vs. M2 and M7 vs. M8): SPO11, TOP6BL, IHO1, REC114, MEI1, MRE11, RAD50, NBN, EXO1, MSH4, HFM1, RNF212, BLM, MUS81, and MLH3 (**Fig. 1*B***). Several of these proteins were identified in earlier studies in other lineages: RAD50 and MRE11 homologs in *Drosophila* (11); MLH1, MRE11B, MUS81-1, and SPO11-2 in maize (13); and IHO1, MSH4, MRE11, and NBS1 in mammals (14). There was overall some, but not perfect, consistency in the ranking of the proteins according to the average D_N_/D_S_ ratio even when signatures of positive selection did not rise to the level of significance in all lineages (**Fig. 1*B***). Taken together, these findings with key recombination proteins corroborate and extend prior reports of signatures of positive selection in rapidly evolving meiotic proteins.

### Signatures of positive selection are enriched in a disordered region of MLH3

MLH3 showed significant evidence of positive selection in all three vertebrate lineages (p < 0.05 for M8 vs. M7; **Dataset S2**), so we focused on this protein to explore the relationship of its domain architecture to its selection signatures. MLH3 is required for crossover-specific resolution of recombination precursors (25, 26) and loss of MLH3 leads to infertility in mice (27). The protein consists of a structured N-terminal domain containing an ATPase head and a transducer domain, as well as a structured C-terminal domain that is required for dimerization with MLH1. The two structured domains are connected by a long intrinsically disordered linker domain (**Fig. 1*C***).

To see where residues that are predicted to be evolving under positive selection are located, we used posterior mean D_N_/D_S_ ratios (from here on: post. mean D_N_/D_S_) and the posterior probabilities that D_N_/D_S_ is greater than 1 generated under Model 8 with Bayes empirical Bayes (**Fig. 1*C*-*D***). Amino acid residues with a posterior probability >0.9 (post. probability, highlighted in pink in **Fig. 1*C***), i.e., those that have the strongest support for being under positive selection, are all within the intrinsically disordered region of the protein. This region is also under less purifying selection in all lineages, as judged by higher post. mean D_N_/D_S_ estimates, although no individual residues reached significance in primates or yeasts (**Fig. 1*C*-*D***).

We therefore tested whether estimating D_N_/D_S_ ratios separately for the structured and intrinsically disordered regions improves detection of selection signatures, using codeml fixed-site models (**Dataset S3**). In this analysis, the linker region is under significantly less purifying selection than the structured domains in all lineages, even in yeast where there is not overall a significant signature of positive selection. MLH1, which shares the same domain structure, showed a similar pattern of less purifying selection in its linker domain (**Dataset S3**). Integrating the post. mean D_N_/D_S_ estimates with Alphafold structural models (in which the linker appears as a disordered ribbon orbiting the structured core) allows direct visualization of these connections between selection rates and domain organization (**Fig. 1*E***).

One potential issue with disordered regions could be poor alignment quality, resulting in erroneously elevated D_N_/D_S_ ratios. To mitigate this concern, we examined MLH3 and MLH1 sequence variation within a single species using mouse population genomic data (28). Since these sequences are close to identical, alignment errors are unlikely. We used DNAsp (29) to analyze intra- and interspecies variation in the linker region and the structured N-and C-terminal domains. In both MLH1 and MLH3, the linker shows more intraspecies variation (**Fig. 1*F***), suggesting that the enrichment of linker sequence divergence within lineages is not (solely) a consequence of alignment ambiguity. In addition to sequence variation, the MLH3 linker also shows length variation between widely diverged species (30) (**Fig. 1*G***). Interestingly, a large part of the linker in mouse (and other species) is encoded by one very long first exon.

Overall, MLH3 provides an instructive example of significant evidence for recurrent positive selection particularly enriched in an intrinsically disordered region. Similar enrichment in intrinsically disordered regions was observed for other proteins in our dataset, including pro-crossover factor HFM1 or the non-crossover-promoting helicase BLM (***SI Appendix* Fig. S2*A*-*D***).

### Evaluating sequence divergence against a backdrop of structural constraints

Disordered regions are, by definition, subject to low structural constraint, allowing many of their residues to vary without negatively impacting protein function. Thus, much if not all the variation in the disordered regions of MLH3, HFM1, and BLM could be due simply to less purifying selection because of less structural constraint (31). To explore this possibility more widely, we asked how the AlphaFold “predicted local distance difference test” score (pLDDT) relates to D_N_/ D_S_ for each residue. A low pLDDT (< 50) can be used as a predictor of disorder (20) (**Fig. 2*A***). When grouped according to this threshold, residues in presumptively intrinsically disordered regions indeed showed higher D_N_/D_S_ estimates on average across all genes. This is true across all lineages and within each lineage considered individually (**Fig. 2*B* and *SI Appendix* Fig. S2*E***).

**Figure 2.**
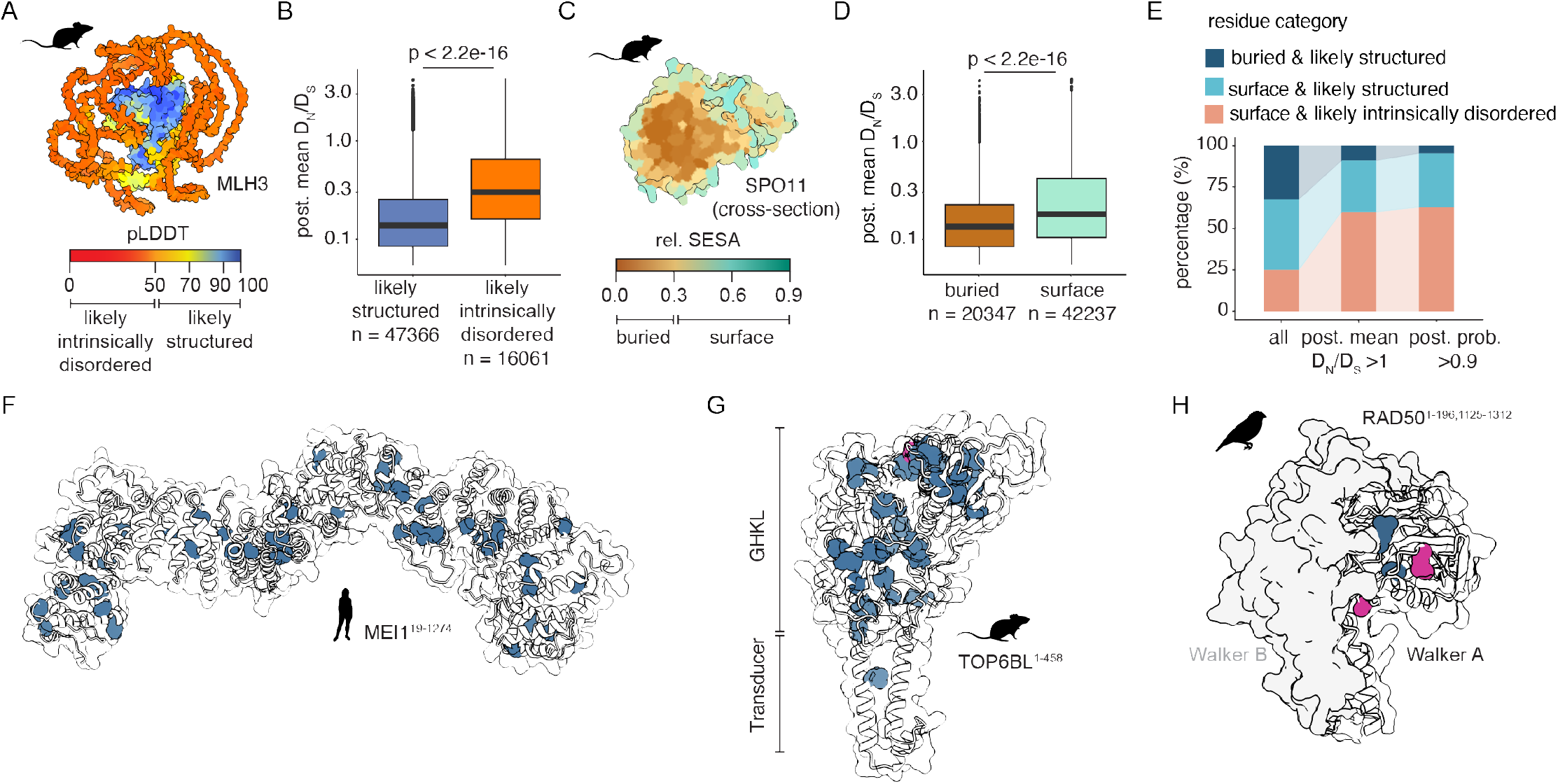
Structural constraints influence selection rates. (*A*) AlphaFold model of mouse MLH3 (from **Fig. 1*E***) colored by prediction confidence score pLDDT. (*B*) Post. mean D_N_/D_S_ values for residues categorized by pLDDT as likely structured or likely intrinsically disordered. The p value is from a t test. Here and throughout, the horizontal lines in boxplots indicate median and interquartile range, whiskers indicate 1.5 times the interquartile range, and outliers are shown as individual points. (*C*) A cross-section of mouse SPO11 AlphaFold structure, colored by the relative solvent exposed surface area (rel. SESA). (*D*) Post. mean D_N_/D_S_ values for residues categorized by rel. SESA as buried or surface. (*E*) Percentages of structured or intrinsically disordered residues in the complete dataset (all residue, all proteins, all lineages) or among only the residues that show relaxed purifying selection (post. mean D_N_/D_S_ > 1) or positive selection (post. probability > 0.9). (*F*-*H*) Transparent surface and ribbon representations of Alphafold models of human MEI1 (*F*), mouse TOP6BL (*G*), and zebra finch RAD50 (*H*). Residues that show relaxation from purifying selection (post. mean D_N_/D_S_ > 1) despite strong structural constraints (buried & likely structured) are in blue (shown as spheres for better visibility, RAD50 residues R3 and S53 in panel *H*); residues in pink (RAD50 M94 and A149 in panel *H*) are also statistically significant (post. probability > 0.9).

By similar reasoning, buried residues are expected to experience stronger structural constraints than surface residues (32). As a measure for an amino acid’s location, we used the relative solvent exposed surface area (rel. SESA) for each residue (see ***SI Appendix, Materials and Methods***) (**Fig. 2*C***). When residues were grouped into buried (rel. SESA < 0.3) and surface categories (rel. SESA >= 0.3), surface residues showed higher D_N_/D_S_ estimates on average, as expected (**Fig. 2*D* and *SI Appendix* Fig. S2*F***).

In the complete dataset, 29.3% of residues were classified as both buried and likely structured (**Fig. 2*E***). By contrast, residues in this category were a smaller fraction of those residues that showed evidence of relaxed selection (8.5% for those with post. mean D_N_/D_S_ > 1) or that more stringently showed evidence of positive selection (4.6% for those with post. probability > 0.9) (**Fig. 2*E***). Conversely, residues classified as surface-exposed and likely intrinsically disordered were overrepresented among residues showing evidence of relaxed or positive selection. Despite this global tendency, we categorized substantial numbers of residues as both buried and likely structured, but that also showed signatures consistent with either relaxed or positive selection (e.g., n = 280 residues with post. mean D_N_/D_S_ > 1; n = 9 residues with a post. probability > 0.9). We consider here several illustrative examples.

The proteins with the most such residues are MEI1 and TOP6BL, which were also among the fastest evolving proteins in our dataset. Although its specific biochemical function remains unclear, MEI1 is required for meiotic DSB formation in mice and *A. thaliana* (33, 34) and mutations in human *MEI1* are associated with infertility (35). Nearly all of MEI1 is predicted to be structured, containing Armadillo-like folds, but 46 residues with a post. mean D_N_/D_S_ > 1 are distributed throughout the core of the structure (**Fig. 2*F***).

TOP6BL is a direct binding partner of SPO11 related to the GHKL-family ATPase subunit of archaeal topoisomerase VI (34, 36). When present, TOP6BL orthologs are essential for DSB formation in every organism tested (10). It has multiple residues that are buried and structured and that also show evidence of positive selection (n = 35 in the rodent dataset and n = 19 in the primate dataset). In rodents, most of these residues are in and around the degenerate GHKL (1-172) as well as at the top of the transducer domain (196-450) (**Fig. 2*G***). These regions overall show more sequence variation than the lower part of the transducer domain, which mediates the interaction with SPO11 (***SI Appendix* Fig. S2*G***). This variation is interesting because the GHKL domain has been lost independently in several lineages (*Drosophila, Schizosaccharomyces, Saccharomyces*) (37, 38).

Remarkably, of the nine residues in the full dataset that scored as buried and structured but also showed significant evidence of positive selection (post. probability >0.9), four are in RAD50 in the bird lineage. RAD50 homologs are universally present in eukaryotes, prokaryotes, and archaea. In eukaryotes, RAD50 is a component of the MRX/N complex and has roles in the early DNA damage response, DSB repair, DSB end resection, and (in some organisms) meiotic DSB formation (39, 40). Two of those four residues (M94 and A149) in RAD50 locate to the N-terminal ATPase domain (Walker A motif), while the other two residues (A332, R788) are in the central flexible coiled coil domain (**Fig. 2*H*, *SI Appendix* Fig. S2*H***). The Walker A domain also contains two additional residues (R3 and S53) that are classified as buried and likely structured and show post. mean D_N_/D_S_ >1 without reaching significance. Components of the MRX/N complex were previously flagged to be under positive selection in mammals and *Drosophila* (11, 14), but it was unclear which protein domains were involved. We conclude that careful evaluation of structural constraints can inform detailed PAML analyses and highlight unexpected sequence variation.

### Structural and functional constraints in protein-protein interaction surfaces

We also evaluated sequence variation within the context of constraints imposed by protein-protein contacts. Similar to structured and buried regions, protein surfaces that mediate protein-protein interactions were found to experience increased purifying selection (41). Our dataset contains four multiprotein complexes: dimers of the SPO11 core complex (the core complex comprises SPO11-TOP6BL heterodimers in vertebrates, or heterotetramers of Spo11, Rec102, Rec104, and Ski8 in yeast), the MRX/N complex (MRE11 and RAD50 plus NBN or yeast Xrs2), MutSγ (MSH4-MSH5), and MutLγ (MLH1-MLH3). We generated models for each complex using AlphaFold3 (42) (**Fig. 3*A***) then identified interfacial contact residues, defined as those that are solvent accessible for a protein in isolation but buried within the complex (**Fig. 3*B*, *see SI Appendix, Materials and Methods***). On average across all complexes in the full dataset, contact residues show significantly lower D_N_/D_S_ than other surface residues. In fact, their average is comparable to constitutively buried residues (**Fig. 3*C*)**.

**Figure 3.**
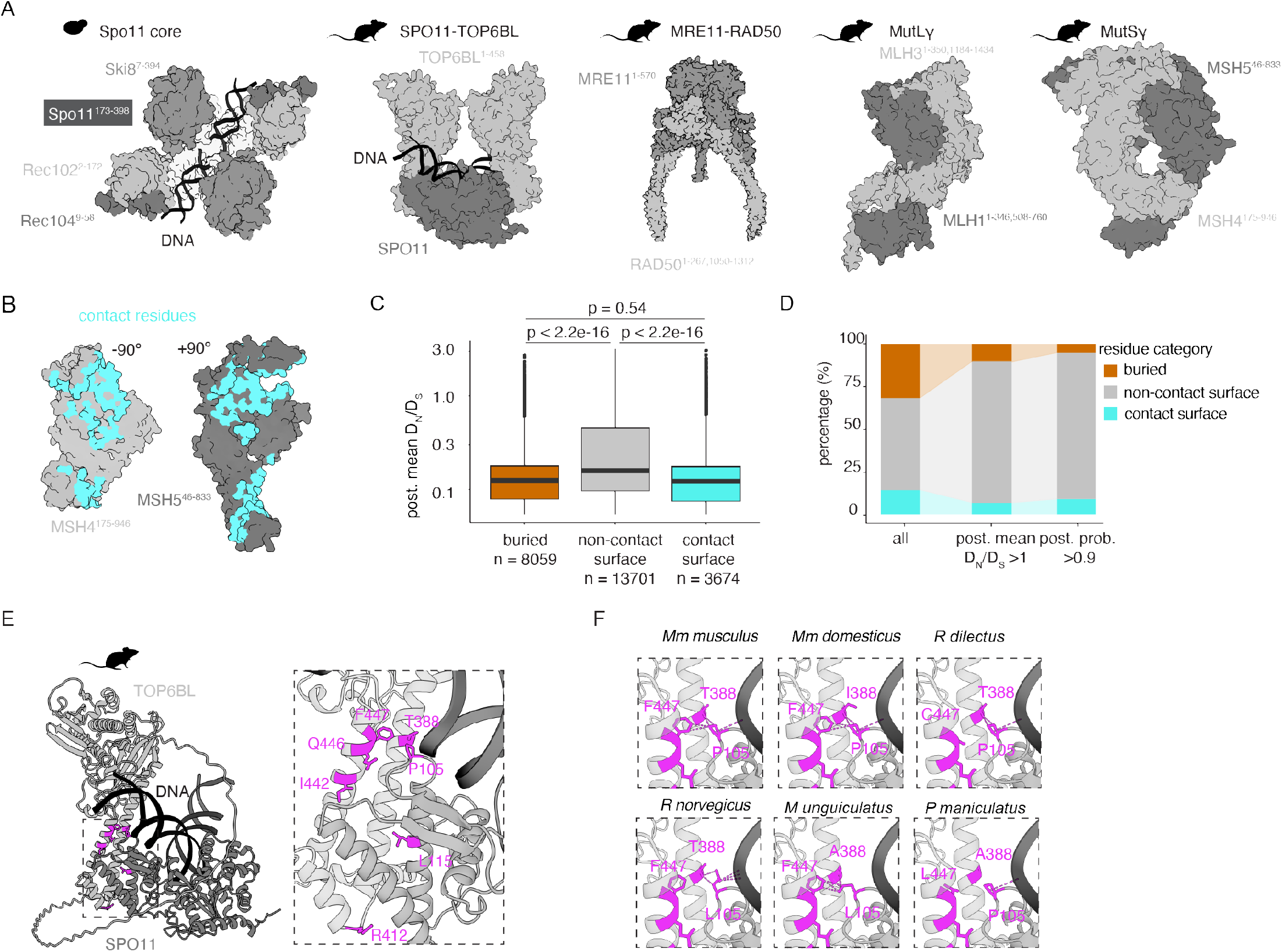
Purifying selection associated with protein-protein interaction surfaces. (*A*) Examples of protein complex structures from cryo-EM or AlphaFold3 prediction. Disordered regions are omitted for clarity. (*B*) MSH4-MSH5 contact residues (cyan) in mouse MutSγ. The proteins are shown as in panel *A*, but each subunit is rotated 90° left or right to reveal the interaction surfaces. (*C*) Post. mean D_N_/D_S_ values for residues characterized as either buried, noncontact surface, or contact surface. The p value is from a t test. (*D*) Percentages of buried, noncontact surface, or surface residues in the complete dataset or among only the residues that show relaxed purifying selection (post. mean D_N_/D_S_ > 1) or positive selection (post. probability > 0.9). (*E*)Overview (left) and detailed view (right) of the interaction between mouse SPO11 and TOP6BL. Only one dimer of SPO11–TOP6BL is shown for simplicity. Interfacial residues with post. mean D_N_/D_S_ > 1 are highlighted in magenta. (*F*) Predicted interaction of SPO11-P105 with TOP6BL-T388 and F447 in *M. musculus musculus* (pink dashed lines, Van der Waals overlap ≥ −0.4 Å). Six distinct amino acid combinations in these three residues are present in the rodent lineage. Four combinations are predicted to partially or completely abolish this interaction when using the swapaa command in ChimeraX. SPO11-P105 is also predicted to interact with the DNA backbone (shown in black).

Thus, as expected, direct protein-protein contacts provide additional structural constraints that we can leverage to give context for interpreting sequence variation. Despite these constraints, and as with constitutively buried residues described in the preceding section, we again identified multiple contact residues displaying indices of relaxed purifying selection (n = 96 residues with post. mean D_N_/D_S_ > 1), out of which 9 residues showed significant signs of positive selection (n = 9 with post. probability > 0.9). The nine significant residues were found in interacting regions of MRE11, RAD50, TOP6BL, MSH4, and MSH5 in different lineages (**Fig. 3*D*, *Datasets S*4-5**).

We consider here the mouse SPO11 core complex as an example. We identified two interfacial residues in mouse SPO11 (P105, L115) with post. mean D_N_/D_S_ > 1, and six in TOP6BL (T388, F447, Q446, I442, R412). Four of the TOP6BL residues (T388, I442, Q446, and F447) cluster in the center of the transducer domain in a region that interacts with loops in the winged-helix domain of SPO11 (**Fig. 3*E***). Interestingly, two of these residues (TOP6BL-T388 and -F447) are predicted to interact directly with one of the variable SPO11 residues (P105) (**Fig. 3*F***). Furthermore, the amino acid combinations present in several other rodent species are predicted to alter or disrupt the interaction (**Fig. 3*F***). A fifth variable TOP6BL residue (R412) and the second SPO11 residue (L115) do not contact one another directly, but both are part of a common contact surface comprising an alpha-helical bundle that is critical for the SPO11-TOP6BL interaction (**Fig. 3*E***) (37, 38, 43, 44). The SPO11-SPO11 dimeric interface also showed three residues with post. mean D_N_/D_S_ > 1 (T195, S197 and L375) (***SI Appendix* Fig. S3*A***). For the same interface in other lineages, primate TOP6BL T388 is conserved as A387, F447 is conserved as F446, Q446 is conserved as Q445, I442 is conserved as I441, and R412 is conserved as R411. For SPO11, P105 and L115 are conserved within primates and birds as well. Within primates two other residues in the vicinity in SPO11 (Q109, S112) show post. mean D_N_/D_S_ > 1 and are also in contact with TOP6BL. TOP6BL itself is conserved at the SPO11 contact surface. In summary, interaction surfaces of protein-protein complexes provide additional structural constraints that can be considered whenever such information is available.

### Leveraging paralogous proteins to highlight otherwise cryptic sequence variation

Proteins that have different biochemical roles may experience different mixes of selection pressures. For example, proteins (or domains within proteins) that have enzymatic activity may be more constrained on average than proteins that serve solely as structural binding partners. Because the variable backdrop of purifying selection may obscure signs of positive selection, we reasoned that it would be advantageous to compare paralogous proteins with similar structures and biochemical properties, but that function in different cellular contexts.

A previous study in mammals found that paralogs that are largely or exclusively meiosis-specific tend to diverge faster (measured as the change in amino acid identity vs. time from last common ancestor) than their counterparts (referred to below as “generic” for simplicity) that have functions outside of meiosis (45). We extended these analyses to additional paralog pairs in plants and budding yeast, finding the same conclusion to apply in nearly all cases (**Fig. 4*A*, *SI Appendix* Fig. S4*A***). Given the broad reproducibility of this trend, we incorporated paralog comparisons into our analyses.

**Figure 4.**
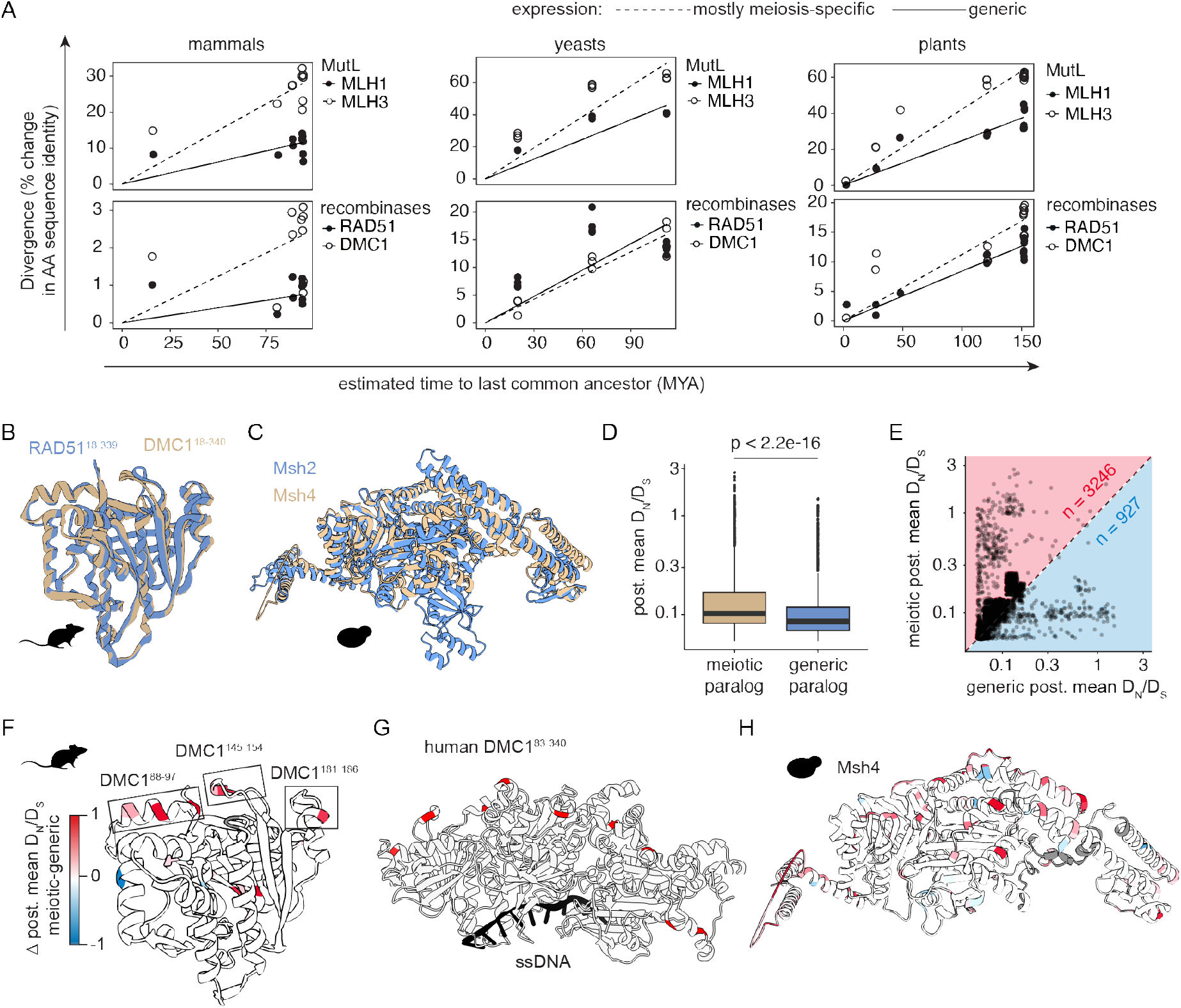
Paralog comparisons reveal subtle sequence divergence. (*A*)Pairwise divergence for selected paralogous protein pairs. Linear regression lines generated in R. Solid circles/ lines: generic expression, open circles/dashed lines: mostly meiosis-specific expression. Analyzed species are *Saccharomyces arboricola, S. kudriavzevii, S. cerevisiae, S. eubayanus, Nauvomozyma castellii*, and *Kluyveromyces lactis* for yeasts, *Arabidopsis thaliana, Brassica napus, B. rapa, Nicotiana tabacum, Oryza sativa* and *Zea mays*, for plants, and *Homo sapiens, Mus musculus, Rattus norvegicus, Bos taurus*, and *Canis lupus* for mammals. (*B*-*C*) Pairwise structural alignment of *M. musculus* RAD51 (blue) DMC1 (brown) in (*B*) and budding yeast Msh2 (blue) and Msh4 (brown) (*C*). (*D*) Post. mean D_N_/ D_S_ values for all residues in the dataset that could be structurally matched between paralogous proteins. The p value is from a t test. (*E*) Scatter plot showing post. mean D_N_/D_S_ values for the same data as in panel *D*. For more residues, the post. mean D_N_/D_S_ is higher in the meiotic paralog (n=3246, residues in the red triangle), but 927 residues have higher post. mean D_N_/D_S_ in the generic paralog (residues in the blue triangle). (*F*) Mouse DMC1, colored by the difference (Δ) in post. mean D_N_/D_S_ between structurally matched residues. Red color indicates higher post. mean D_N_/D_S_ in DMC1 while blue color indicates more variability in RAD51. Three loops that show sequence variation specifically in DMC1 are highlighted. (*G*) The residues of interest identified in panel *F* highlighted in red on a crystal structure of human DMC1 bound to single stranded DNA (PDB: 7C9C). (*H*) Yeast Msh4, colored as in panel *F* based on comparison with Msh2.

In a first step, we tested if the strictly meiotic proteins DMC1 (meiosis-specific strand exchange protein) and MSH4 (a subunit of the MutSg complex) show higher D_N_/D_S_ ratios than their generic paralogs RAD51 and MSH2 (subunit of MutSa and MutSb (46)), respectively. As predicted, codeml fixed-site models improve when we allow for different D_N_/D_S_ ratios between the two genes in each meiotic vs. generic paralog pair in all lineages except for RAD51 vs. DMC1 in the bird lineage (**Dataset S6**). In addition, the meiotic paralog shows a higher average D_N_/D_S_ value in all cases except yeast Rad51 and Dmc1 (**Dataset S6**).

Armed with this information, we then set out to directly compare structurally matched residues between paralogs, reasoning that such a comparison would help control for the differences in selection pressures experienced by different types of proteins and by residues residing in different biochemical/functional contexts. To do this, we generated pairwise structure alignments for RAD51 vs. DMC1 (**Fig. 4*B***) and MSH2 vs. MSH4 (**Fig. 4*C***), yielding 4267 residues that could be matched with high confidence between the paralogs in the four lineages. In agreement with the gene-level models, the residues of the meiotic paralogs show a significantly higher post. mean D_N_/D_S_ on average (**Fig. 4*D***). When D_N_/D_S_ values for individual pairs of structurally matched residues are examined, 3246 show higher D_N_/D_S_ in the meiotic paralog and only 927 are higher in the generic paralog (**Fig. 4*E***).

We further mapped the differences between meiotic and generic D_N_/D_S_ values onto the Alphafold structures (**Fig. 4*F*-*H***). This analysis highlighted a set of rapidly diverging residues located near one another on a surface of rodent DMC1 (**Fig. 4*F***). Closer inspection showed that one residue (R94), shows post. mean D_N_/D_S_ > 1, and two more (T148 and E182) are just below that threshold (D_N_/D_S_ = 0.943 and 0.891 respectively) (***SI Appendix* Fig. S4*B*-*C***). In contrast, the equivalent residues are widely conserved in DMC1 in the other lineages. This surface is opposite the ssDNA-binding region, such that these variable residues spiral around the nucleoprotein filament formed by cooperative assembly of DMC1 on ssDNA (**Fig. 4*G***).

As a second example we analyzed yeast Msh4 (**Fig. 4*H***). MSH4 showed significant signs of positive selection in the primate and rodent lineages but not in yeast (***SI Appendix* Fig. S4*D*-*G* Dataset S2**). Nevertheless, yeast Msh4 contains many residues that are specifically under less purifying selection in Msh4 than in Msh2, but unlike in mouse DMC1, they are more broadly distributed throughout the structure (**Fig. 4*H***). We conclude that comparative analysis of meiotic and generic paralogs provides a useful framework to test for potential functional variation in more conserved or structurally constrained proteins.

### Functional impact of sequence variation in budding yeast MSH4

Yeast provides a straightforward experimental system for gene-swap experiments to test the functional relevance of sequence variation. Based on the paralog comparison in the yeast clade, we sought to test whether the sequence variation in yeast Msh4 affects function, despite not rising to the level of statistical support for positive selection. We performed cross-complementation experiments using FLAG-tagged *MSH4* gene sequences from *Saccharomyces* species *S. paradoxus, S. bayanus* and *S. arboricola* expressed in *S. cerevisiae* under the control of the endogenous *MSH4* promoter (**Fig. 5*A***). Immunoblots confirmed expression and stability of the proteins in synchronously sporulating cultures (***SI Appendix* Fig. S5*A***).

**Figure 5.**
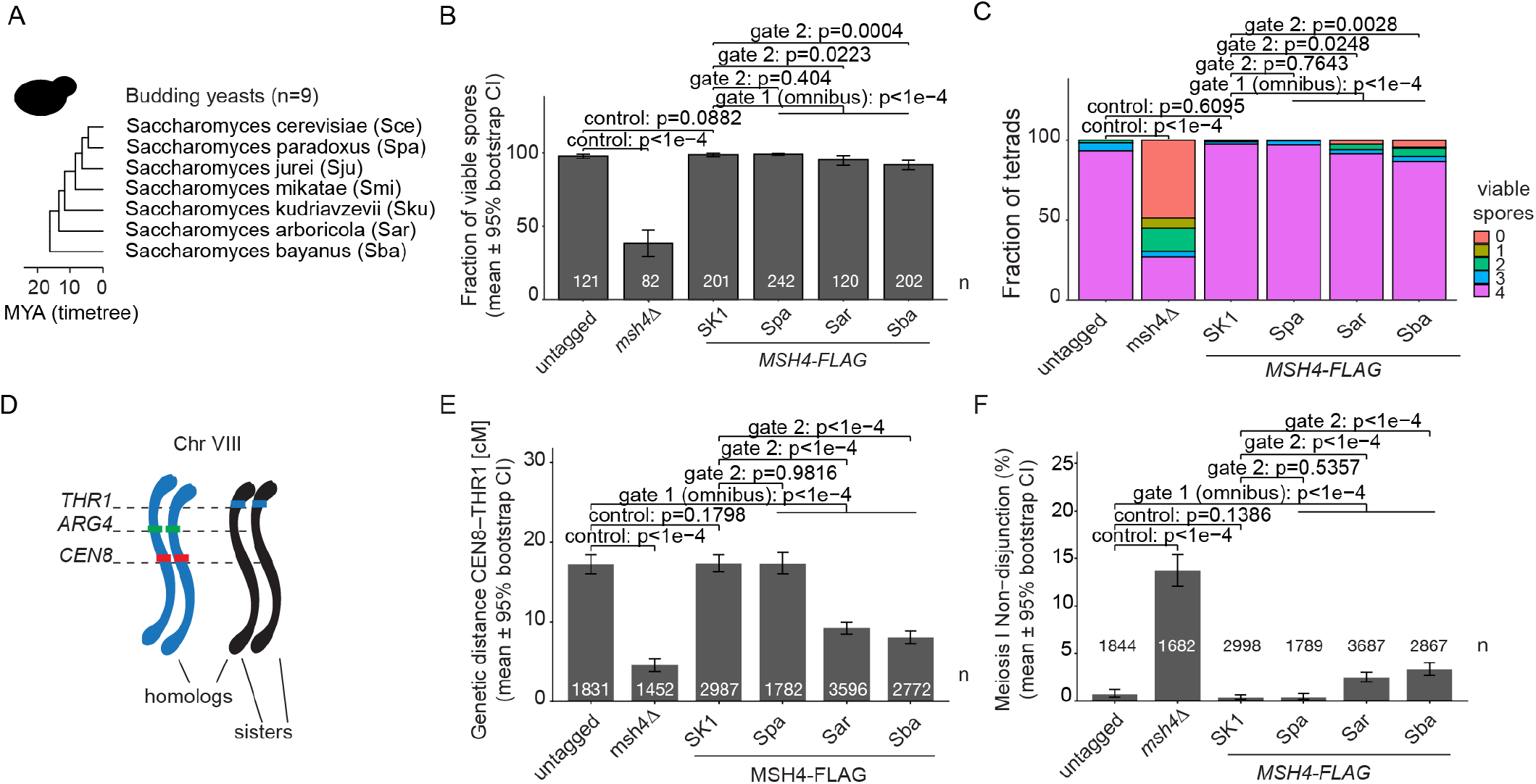
Cross-complementation assays with budding yeast Msh4. (*A*) Endogenous *MSH4* in *S. cerevisiae* was replaced with FLAG-tagged versions of *MSH4* from diverged *Saccharomyces* species (bold). (*B*) Spore viability. Bars show the mean and error bars indicate 95 % bootstrap confidence intervals. The total number of tetrads dissected is indicated for each genotype. P-values are indicated for comparisons using beta-binomial generalized linear model testing. Holm’s adjusted values are shown for gate 2 tests. (*C*) To detect signatures of MI-NDJ events, the number of viable spores was scored for each tetrad. Stacked bar plots show the mean number of viable spores per tetrad for the same tetrad data as in panel *B*. P-values are indicated for comparisons as in panel *B*. (*D*) Spore-autonomous fluorescent constructs allow the scoring of crossing over in the *CEN8–THR1* interval. (*E*) Genetic distance data for the *CEN8–THR1* interval was calculated using the Perkin’s equation. Bars indicate mean genetic distances and error bars represent 95% bootstrap confidence intervals. The total number of tetrads for each genotype is indicated excluding NDJ events. Fisher’s test p-values are indicated for selected comparisons. Holm’s adjusted values are shown for gate 2 tests. (*F*) MI-NDJ was calculated from the same tetrad data as in panel *E*. Bars show the mean, error bars indicate 95% Wilson binomial confidence intervals. Fisher test p-values are indicated for the indicated comparisons. Holm’s adjusted values are shown for gate 2 tests and the total number of tetrads analyzed is indicated.

Using tetrad dissection, we found that *MSH4* from divergent species supported better spore viability than a *msh4Δ* null, but to different extents (**Fig. 5*B*-*C***). While reintegration of *S. cerevisiae MSH4* or complementation with *S. paradoxus MSH4* was indistinguishable from the wild-type control, complementation with *MSH4* from *S. arboricola* and *S. bayanus* yielded a small decrease in viability (**Fig. 5*B***).

To test whether this decrease is statistically significant we used a hierarchical testing strategy: we first assessed genotype effects using a beta-binomial generalized linear model and tested whether any heterologous *MSH4* allele differed from the reintegration control (omnibus test) in overall viability. Since this test demonstrated a significant difference (p = 5.84E-06), we then tested each heterolog individually against the reintegration control using one-sided tests with Holm-Bonferroni adjustment. Both *S. arboricola* and *S. bayanus MSH4* differed significantly from the control (adjusted p = 0.0223 and p = 0.0004, respectively), so we then performed stepwise contrasts (*S. arboricola* vs. *S. paradoxus*; *S. bayanus* vs. *S. arboricola*) to assess evolutionary ordering. *S. arboricola MSH4* performed significantly worse than *S. paradoxus MSH4* (adjusted p = 0.0246). *S. bayanus* gave a further decrease from *S. arboricola*, but this was not significant after correction (adjusted p = 0.0828) (**Fig. 5*B*, Dataset S7**).

Given the role of Msh4 in recombination, we asked whether the decrease in spore viability is due to a disproportionate increase in the frequencies of tetrads with two or zero viable spores, which is the pattern predicted for chromosome missegregation during the first meiotic division caused by recombination defects (47). Analyzing the tetrad data for this signature, we found a significant increase in tetrads with two or four dead spores in the strains complemented with *MSH4* from *S. arboricola* (adjusted p = 0.0248 compared to the reintegration control) or *S. bayanus* (adjusted p = 0.0028) (**Fig. 5*C*, Dataset S7**).

To directly test for defects in recombination, we measured crossing over within the *CEN8–THR1* interval on chromosome VIII using genetic markers that provide spore-autonomous expression of fluorescent proteins (48) (**Fig. 5*D*, *SI Appendix* Fig. S5*C***). We employed a similar hierarchical testing strategy as above but using Fisher’s exact test and bootstrap analysis to assess differences in recombination (**Dataset S8**). The omnibus test showed that at least one heterologous strain differed in the genetic distance across the whole *CEN8–THR1* interval (Fisher’s adjusted p = 4.77E-83). Individual comparisons showed a significant reduction in genetic distance upon complementation with *MSH4* from *S. arboricola* (adjusted p = 1.52E-45) or *S. bayanus* (adjusted p = 3.48E-52) but not *S. paradoxus* (adjusted p = 0.982) (**Fig. 5*E***). *S. bayanus MSH4* performed worse than *S. arboricola* (adjusted p = 5.88E-05; **Dataset S8**), consistent with expectation for evolutionary ordering. Complementary bootstrap tests confirmed these findings (**Dataset S8**).

The fluorescent spore data also provide a means to assess meiosis I nondisjunction (MI-NDJ) of chromosome VIII (48). The strains with *S. arboricola* or *S. bayanus MSH4* showed a significant increase in MINDJ events compared to the reintegration control (Fisher’s adjusted p = 1.29E-13 and p = 9.05E-19, respectively) (**Fig. 5*F*, Dataset S9**). Further consistent with expectation based on evolutionary distances from *S. cerevisiae*, the cross-complementation phenotype was worse for *S. bayanus* than *S. arboricola* (one-sided Fisher’s adjusted p = 0.0248) and *S. arboricola* was worse than *S. paradoxus* (one-sided Fisher’s adjusted p = 2.32E-09) (**Dataset S9**). These findings, which are consistent with the tetrad-level spore viability signatures, show that the recombination defects caused by heterologous Msh4 expression result in more frequent chromosome segregation errors and reduced fertility.

Since Msh4 works as part of a heterodimer with Msh5, we considered the possibility that the observed defects might be due to incompatibility between the heterologous Msh4 and *S. cerevisiae* Msh5. Within-strain sequence variation can cause such incompatibility for MutLα complexes (Mlh1–Pms1) in *S. cerevisiae* (49, 50). If true in our system, it predicts that replacing the *S. cerevisiae* Msh5 with the species-matched Msh5 would improve the ability of heterologous Msh4 to cross-complement. Alternatively, because the *S. cerevisiae* Msh5 likely retains most or all its own specific interactions and behaviors, replacing it with the heterologous Msh5 proteins might exacerbate defects of the heterologous Msh4 proteins.

Immunoblots of myc-tagged versions of Msh5 heterologs from *S. arboricola* and *S. bayanus* confirmed protein stability across meiotic time courses (***SI Appendix* Fig. S5*B***). To assess spore viability and recombination, we introduced untagged versions of Msh5 into the cognate cross-complemented Msh4 strains (***SI Appendix* Fig. S5*D*-*G***). Strikingly, partnering *S. bayanus MSH4* and *MSH5* substantially decreased overall spore viability and crossing over and increased MI-NDJ of chromosome VIII compared to *S. bayanus MSH4* alone (***SI Appendix* Fig. S5*D*-*G***; results of statistical tests are in **Datasets S7-9**). These findings appear to match the second scenario outlined above. In contrast, partnering *S. arboricola MSH4* and *MSH5* did not consistently improve or worsen meiotic outcomes compared to *S. arboricola MSH4* alone (***SI Appendix* Fig. S5*D*-*G***).

### Lineage specificity of rapid sequence change

Previous studies found that signs of positive selection were often restricted to specific lineages (11, 13, 14). Our whole-gene evolutionary analysis similarly showed different levels of purifying, relaxed, or positive selection between lineages for several proteins (**Fig. 1*C***), and the sequence variation discussed above for the SPO11–TOP6BL interface (**Fig. 3*F***) and loop elements in DMC1 (**Fig. 4*F***) is specific for rodents. We examine here several additional examples.

As described above, the N-terminal ATPase domain of RAD50 in birds has two buried residues with post. mean D_N_/D_S_ > 1 (zebrafinch R3 and S53) and two residues with post. probability > 0.9 (M94 and A149; **Fig. 2*H***). In addition, seven surface residues in this domain (V17, S53, V93, L101, G106, T108, T121) also have post. mean D_N_/D_S_ > 1 in birds, with post. probability > 0.9 for residues 101 and 108 (**Fig. 6*A,B***). This domain also displays substantial variability in rodents but affecting mostly different residues: mouse M93 and A95 have post. mean D_N_/D_S_ > 1 and M121 and K126 additionally have post. probability > 0.9 (**Fig. 6*A,C***). Additionally, we identified two residues (T1193, S1309) in the mouse Walker B motif with D_N_/D_S_ > 1. By contrast, the whole domain is highly conserved in both primate and yeast lineages (post. mean D_N_/D_S_ < 1 for all residues; **Fig. 6A and *SI Appendix* Fig. S6A**).

**Figure 6.**
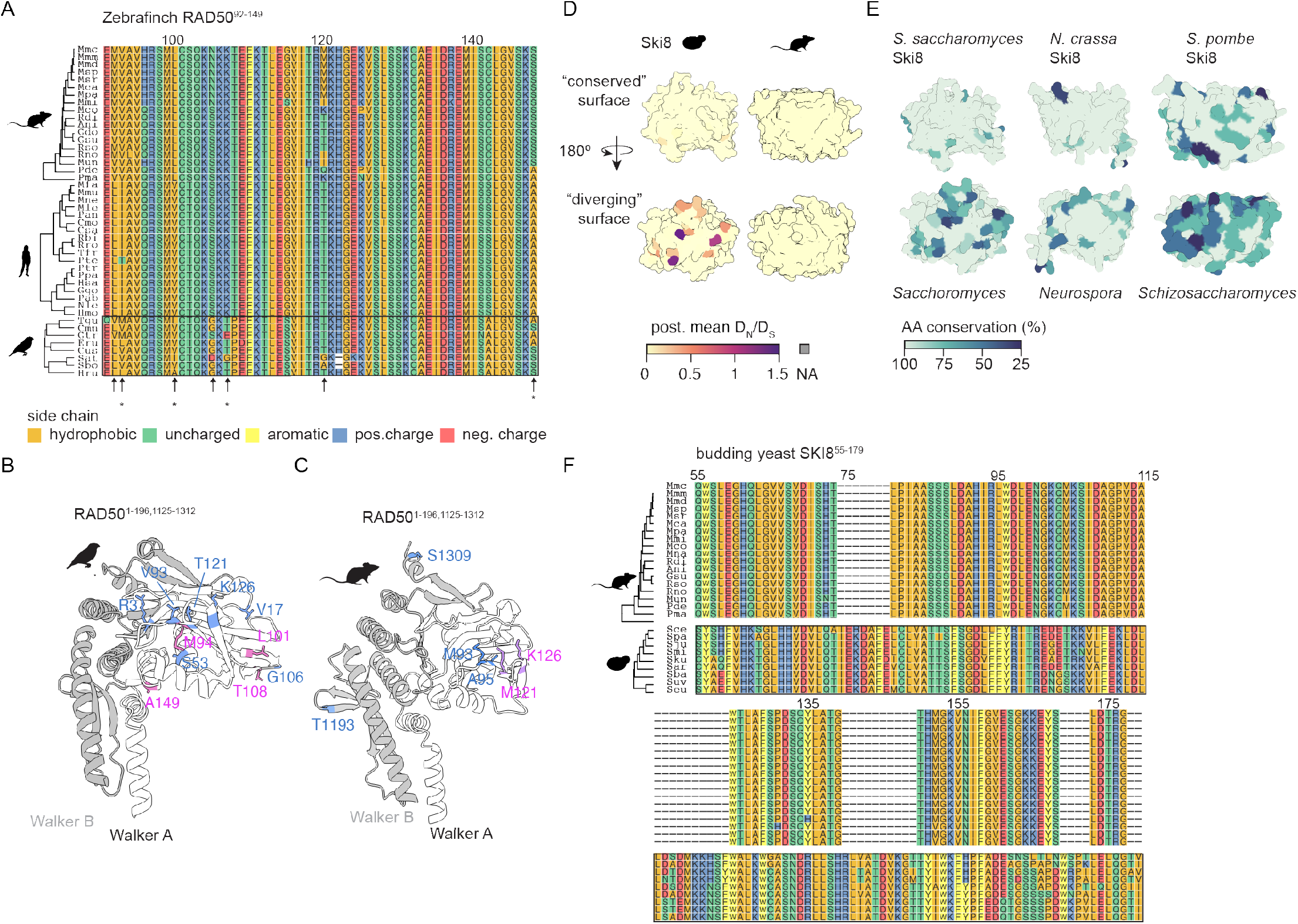
Lineage specificity of sequence variation. (*A*) Partial alignment of the Walker A domain of RAD50. Amino acids are colored according to biochemical side chain properties (Hydrophobic: I, A, V, L, P, G, M; uncharged: Q, T, C, N, S; aromatic: Y, F, W; positive charge: R, H, K; negative charge: D, E). Black rectangle indicates the lineage in which we initially observed relaxation from purifying selection. Arrows indicate residues with post. mean D_N_/D_S_ > 1 and asterisks indicate residues that also have post. probability > 0.9. (*B*-*C*) RAD50 ATPase domains of zebra finch (*B*) and mouse (*C*). Blue residues show post. mean D_N_/D_S_ > 1; and pink residues also show post. probability > 0.9. (*D*) Post. mean D_N_/D_S_ mapped onto the Alphafold model of budding yeast Ski8 (left) and mouse WDR61 (right). Surface representations are matched between the species: N-terminal half above (more conserved in yeast), C-terminal half below (more diverged in yeast). (*E*) Alphafold models of *Neurospora crassa* Ski8 (AF-Q7RVN5-F1) and *S. pombe* Ski8 (AF-Q09150-F1) together with *S. cerevisiae* Ski8 from panel *D*. Structures are colored by percent amino acid identity based on multiple sequence alignments (*N. crassa, N. cerealis, N. tetrasperma, N. Africana, N. sublineolata*, and *Sordaria macrospora* for the *Neurospora model*; *S. pombe, S. japonicus, S. cryophilus*, and *S. octosporus* for the *Schizosaccharomyces* model. (*F*) Partial alignment of the N-terminal half of yeast Ski8 with equivalent region of rodent WDR61 (the primate and bird sequences are nearly identical to the rodent sequences)

Ski8 in budding yeast provides another example of lineage-specific sequence variation. Ski8 directly interacts with Spo11 and is essential for meiotic DSB induction (51). This role is conserved in *S. pombe* and *Sordaria macrospora* (52, 53). However, Ski8 is dispensable for meiosis in *A. thaliana* (54)**ÿ**, and it is unclear whether its orthologs have meiotic roles in metazoans or other lineages. Another function of Ski8 is more widely conserved: it forms a complex with Ski2 and Ski3 that influences RNA metabolism by the exosome (55). Ski8/WDR61 was one of the most conserved proteins in our dataset, showing strong purifying selection (**Fig. 1*B***).

Nonetheless, Ski8 in the *Saccharomyces* clade displayed multiple variable residues, most of which were confined to one half of the toroidal structure formed by the protein’s WD40 repeats (**Fig. 6*D*, *SI Appendix* Fig. S6*B*-*C***). The same surface is completely conserved in the rodent lineage (**Fig. 6*C***). In addition to these residues on the protein surface, several buried residues in the same part of the protein showed evidence of relaxed purifying selection as well (post. mean D_N_/D_S_ > 0.4; ***SI Appendix* Fig. S6*B*-*C***). Overall, we found that despite strong purifying selection across the whole protein, the N-terminal half of Ski8 (*S. cerevisiae* residues 12-179) has significantly higher D_N_/D_S_ when using PAML fixed-site models (**Dataset S3**). Notably, the variable half of Ski8 is surface-exposed in both the Spo11 core complex and the Ski2–Ski3–Sk8 complex, whereas the more conserved face interacts directly with Ski2 in the Ski complex (***SI Appendix* Fig. S6*D*-*E***). We also observed surface variability in this half of the protein in other fungal lineages (namely, *Neurospora* and *Schizosaccharomyces*; **Fig. 6*E***). The budding yeast protein also has several insertions of loops between the highly structured beta-sheets (visible as gaps in the other species in the alignment) (**Fig. 6*F***). By contrast, WDR61 is almost perfectly conserved across the whole set of mammalian species, indicating high levels of purifying selection (**Fig. 6*F***). These findings illustrate how comparisons between homologs in different lineages can pinpoint sequence variation that may be functionally relevant.

## Discussion

In this study, we explored how sequence evolution correlates with measures of intrinsic disorder, buried versus surface amino acid side chains, and the interfaces of protein-protein contacts within complexes. As anticipated, residues displaying a greater degree of structural and biochemical constraints were more likely to also display stronger purifying selection, but there were also many exceptions to this general feature. Our results show the value in integrating structural information to contextualize protein sequence changes with respect to domain architecture and biochemical function, and to unmask signs of relaxed or positive selection that may be obscured against a backdrop of purifying selection. AlphaFold greatly expands the information available for this contextualization, which facilitated the development of tools to identify rapidly evolving proteins that incorporate structural details to uncover these signatures (22, 56). We anticipate that structure-informed evolutionary analyses can be extended more broadly to other tools beyond PAML. Furthermore, we anticipate that the strategies explored here will be valuable discovery tools to document evolutionarily relevant sequence changes and to design targeted genetic experiments, including, for example, inter-species complementation tests by swapping adaptive variants or introducing chimeric variants to understand the functional consequences of sites under positive selection (57, 58).

One promising strategy we employed was the direct comparison of paralogous proteins that are matched for structural and functional properties and that have been evolving within the same species (45). Such comparisons provide a controlled like-for-like match of residues that are in a similar structural and biochemical context and are in proteins that perform similar molecular tasks. The approach thus has the potential to highlight sequence variation that is functionally significant even if positive selection is difficult to detect with evolutionary analysis methods that have limited statistical power. This approach should be broadly applicable to any paralog pairs, and it may also extend to comparisons between homologous domains of non-paralogous proteins.

Many proteins lack paralogs within the same species. With these cases, an alternative is to systematically compare between lineages for matched residues of orthologs. As with paralogs, this approach also controls for the backdrop of functional constraints while searching for signatures of positive selection. The lineage-specific examples we document here (the SPO11–TOP6BL interface; loop elements in DMC1; the RAD50 ATPase domain; and a specific surface on Ski8) join results from earlier studies (11, 13, 14). Because each lineage represents a distinct evolutionary trajectory with different histories of selection pressures and adaptations, lineage- or species-specific adaptations in the meiosis and recombination machinery can have crucial reproductive fitness consequences (12, 59). Thus, lineage specificity per se may be a useful hallmark to infer the functional relevance of observed sequence variation, and to highlight candidates for mechanistic follow-up studies.

Our data add to growing evidence that the meiotic recombination machinery, despite its essential functions for fertility, has pervasively experienced rapid evolution including relaxed purifying selection and/ or positive selection (reviewed in (10)). For several proteins, their rapid evolution contributes directly to variation in the recombination process itself. For example, changes in the DNA-binding sequence specificity of the histone methyltransferase PRDM9 cause changes in the distribution of recombination events across the genomes of many vertebrates (60, 61), and variation in amino acid sequence and/or gene copy number of the SUMO E3 ligase RNF212 is associated with variation in global crossover frequencies (62, 63).

In most cases, however, the causes and consequences of rapid evolutionary changes of meiotic recombination proteins are not known. The sequence divergence we observe could be a result of relaxed purifying selection, adaptation to endogenous or exogenous selection pressures, genetic drift, increased mutation rates, or a combination of those. Several specific selection pressures have been proposed to play a role in shaping the recombination machinery: karyotype changes, the evolution of sex chromosomes, genomic conflict or meiotic drivers, and environmental effects including temperature (discussed in (10)). Other factors that could shape meiotic protein evolution in a lineage-restricted manner could include multicellularity vs. unicellularity or sex differences in meiosis and gametogenesis (e.g., oogenesis vs. spermatogenesis).

Interestingly, two of the examples highlighted here have antiviral functions in certain lineages in addition to meiotic recombination functions. Antiviral functions as seen in host-pathogen conflicts are another classic driver of rapid evolution. First, the MRN complex inhibits adenovirus infection in mammalian cells and is in turn targeted for downregulation by a virus-encoded factor (64, 65). Second, the Ski2– Ski3–Ski8 complex reduces copy number and suppresses expression of double-stranded RNA viruses in *S. cerevisiae* (thus the “superkiller” phenotype when Ski proteins are missing because of overexpression of a virus-encoded secreted toxin) (66). It is thus interesting to consider that lineage-specific signatures of rapid evolution of these factors might reflect selection pressures related to host-pathogen conflicts, meiosis roles, or both.

## Materials and Methods

### Sequence curation and phylogenetic analyses using maximum likelihood

Protein coding sequences were collected from public databases or identified in BLAST searches for the genes outlined in **Dataset S1**. Multiple sequence alignments were manually curated and are available on Figshare. Maximum-likelihood analyses were performed with codeml (PAML 4.9) (17), using site models M0, M1, M2, M7 and M8 to estimate D_N_/D_S_ and to test for positive selection by likelihood ratio test. Post. mean D_N_/D_S_ estimates from model M8 (Bayes empirical Bayes method) were mapped onto AlphaFold protein structures in ChimeraX (67) using custom R scripts (Figshare), with a posterior probability threshold of 0.9 used to call significant positive selection. Selection rates were also compared between paralogous proteins and protein domains using PAML fixed-site models (68), with domain boundaries from Interpro (69). Full details, including model comparisons and robustness checks, are provided in ***SI Appendix, Materials and Methods***.

### Analyses of mouse population genomic data

Population genomic data for 23 wild mouse strains from five geographic populations (28) were used to reconstruct strain-specific genomes and extract RefSeq gene sequences. McDonald-Kreitman tests, with *Mus spretus* as an outgroup, were used to calculate the direction of selection (DoS) statistic (DNAsp) (29). Custom scripts are available on Figshare. Full pipeline details are provided in ***SI Appendix, Materials and Methods***.

### Protein structure prediction and visualization

Structural models were obtained from the AlphaFold database (21) or, when unavailable, generated with ColabFold (70). Structures were visualized, and solvent-exposed surface area (SESA) analyzed, in UCSF Chimera (71) or ChimeraX (67). The yeast Spo11 core complex was modeled as a dimer based on the published cryo-EM structure (38); other complexes were predicted using AlphaFold3 (42). Interface residues were classified as buried, surface, or contact surface based on relative SESA and involvement in interchain contacts (72). Interspecies differences at these positions were examined using ChimeraX’s *swapaa* function. Structural divergence between paralogous proteins (RAD51/DMC1, MSH2/MSH4) was assessed by FATCAT alignment (73). Protein domain annotations are from Interpro (69). Full thresholds and parameters are provided in ***SI Appendix, Materials and Methods***.

### Yeast methods

All strains are SK1 derivatives and listed in ***SI Appendix* Table S1**. Spore-autonomous fluorescent reporter strains were as described previously (48). The endogenous *MSH4* locus was replaced with a FLAG-tagged allele by In-Fusion cloning (Takara) and integrated into the wild-type SK1 background. Similarly, the endogenous *MSH5* locus was replaced with a myc-tagged allele. Protein samples were prepared from synchronously sporulating cultures and analyzed by SDS-PAGE and immunoblot, using an anti-FLAG antibody to detect Msh4, an anti-myc antibody to detect Msh5 and an anti-tubulin antibody as a loading control. Full culture and blotting conditions are provided in ***SI Appendix, Materials and Methods***.

Spore viability was determined by tetrad microdissection following sporulation and analyzed using a beta-binomial generalized linear model, with pre-specified contrasts tested in a hierarchical order reflecting the expected evolutionary ordering of heterologous complementation defects. MI-NDJ frequencies were analyzed similarly using logistic regression. Full statistical models and justification for test sidedness are provided in ***SI Appendix, Materials and Methods***.

Spore-autonomous fluorescence recombination assays were performed as described previously (48). Tetrads were scored manually to calculate genetic map distances using the Perkins equation and MINDJ frequencies for chromosome VIII. Genotypes were compared using Fisher’s exact tests, with planned contrasts following the same hierarchical structure used for the viability analysis. Full imaging, scoring and bootstrap procedures are provided in ***SI Appendix, Materials and Methods***.

### Generative AI use

ChatGPT (version 5.2, OpenAI) and Claude 4 Sonnet (Anthropic, San Francisco, CA, USA) were used to assist in developing the statistical framework, writing R scripts, and drafting methods descriptions. Claude Science (Anthropic, San Francisco, CA, USA), running Claude Sonnet 5, was used to execute and independently reproduce the statistical analyses, annotate the R scripts for public distribution, and cross-check reported statistics against the manuscript and response-to-reviewers text for accuracy. After using these tools, the authors reviewed and edited the content for accuracy and take full responsibility for the content of the published article.

## Data, materials, and software availability

Study data are included in the article and/or SI Appendix. Further raw data, sequence alignments and custom scripts are available at Figshare at https://figshare.com/projects/Structure-informed_evolutionary_analysis_of_the_meiotic_recombination_machinery/263293). Individual digital object identifiers (DOIs) are listed in SI Appendix.

## Acknowledgments

We thank Corentin Claeys Bouuaert for plasmids and yeast strains. We thank members of the Keeney laboratory for discussions. MSK core facilities are supported by National Cancer Institute Cancer Center support grant P30 CA08748. Work in the Lai lab was supported by the National Institutes of Health (R01 HD108914 and R01 GM083300). M.A. was supported in part by an EMBO long-term fellowship (ALTF 905-2019) and a postdoctoral award from the SKI Basic Research Innovation Award Initiative and the Dewitt Wallace Basic Science Award Fund. J.V. was supported by a K99/R00 award from the NIH (GM137077). Research in the Keeney lab is supported by NIH grants R35 GM118092 (to S.K.), and R01 HD110120 (to S.K. and D. Patel).

This article is subject to HHMI’s Open Access to Publications policy. HHMI lab heads have previously granted a nonexclusive CC BY 4.0 license to the public and a sublicensable license to HHMI in their research articles. Pursuant to those licenses, the author-accepted manuscript of this article can be made freely available under a CC BY 4.0 license immediately upon publication.

