## Supplementary material for "Structure-informed evolutionary analysis of the meiotic recombination machinery": SI Appendix

**Supporting Information for**
Structure-informed evolutionary analysis of the meiotic recombination machinery

Meret Arter, Jeffrey Vedanayagam, Min Lu, Momar Diop, Kaixian Liu, Eric C. Lai, Scott Keeney

**This PDF file includes:**

Supplemental Materials and Methods
Figures S1 to S6
Table S1
SI References

**Supporting datasets for this manuscript provided as an Excel file:**

**Dataset S1.** Gene list.
**Dataset S2.** PAML results.
**Dataset S3.** PAML domain results.
**Dataset S4.** Interacting residues under positive selection.
**Dataset S5.** Protein-protein complex analyses.
**Dataset S6.** PAML paralog results.
**Dataset S7.** Statistics tetrad viability.
**Dataset S8.** Statistics genetic distance.
**Dataset S9.** Statistics Meiosis I non-disjunction.

**Raw data and scripts available on Figshare:**

**Raw data and scripts:** https://doi.org/10.6084/m9.figshare.30071095.v3
**Sequence alignments and trees:** https://doi.org/10.6084/m9.figshare.30071203.v1
**PAML control files:** https://doi.org/10.6084/m9.figshare.30071092.v1
**PAML results fixed site models:** https://doi.org/10.6084/m9.figshare.30071161.v1
**PAML results pairwise:** https://doi.org/10.6084/m9.figshare.30071164.v1
**PAML results M0, M1, M2, M7, M8:**
https://doi.org/10.6084/m9.figshare.30071167.v1
**Protein structure predictions:** https://doi.org/10.6084/m9.figshare.30071176.v1
**Protein complex predictions:** https://doi.org/10.6084/m9.figshare.30071191.v2
**Attribute files:** https://doi.org/10.6084/m9.figshare.30071209.v1

### Supplemental Materials and Methods

**Sequence curation and phylogenetic analyses using maximum likelihood.** Sequences were collected from NCBI, Ensembl and UniProt databases (74-76). We used the isoform that had been confirmed experimentally, or, in the absence of such information, the longest annotated isoform. Our initial database contained a list of 1415 protein coding sequences covering 91 genes from 4 lineages. Out of those, 240 sequences were re-annotated using blastn because they were either mis-annotated or not previously annotated. Overall, 111 sequences could not be identified, likely due to gaps in the genome assemblies. Additional sequences were excluded from further analyses if they contained large gaps, leading to a total of 259 missing sequences. Because the robustness of PAML parameter estimation is strongly dependent on sequence number we further excluded any gene from a lineage with less than five homologous sequences available. Codon-guided multiple DNA sequence alignments were generated using MUSCLE as implemented in the R muscle package (23). Multiple sequence alignments were then manually curated to remove gaps and alignment errors. Sequence alignments are available on Figshare. To check whether the sequence divergence was comparable between lineages (and thus suitable for PAML analysis) we ran pairwise codeml (model 0) to estimate pairwise synonymous and non-synonymous substitution rates (***SI Appendix*** **Fig. S1*E*)** (17, 77, 78). To test whether synonymous sites in the yeast clade are approaching saturation, we performed substitution saturation analysis using the R package ape: version 5.0 (79). Pairwise distances were calculated using the `ape::dist.dna` function, with the number of transitions, transversions, and the Kimura-two parameter distance per pair identified using the following models: `model = "TS"` for transitions, `model = "TV"` for transversions, and `model = "K80"` for the Kimura two-parameter distance. Our analysis found that third-position transitions show pronounced negative curvature against corrected distance (***SI Appendix*** **Fig. S1*G*)**. This means transitions in the most diverged species pairs already differ at 31.1% of sites, exceeding the 23.2% expected between fully randomized sequences with the same base composition, while transversions remain at 13.8% against an expected 50.0%. Transitions have therefore passed their peak and are declining toward equilibrium, which is a signature of substitution saturation.

**PAML site models.** Phylogenetic analyses using maximum likelihood were performed using codeml from the PAML 4.9 suite (17). Species trees (***SI Appendix*** **Fig. S1*A-D***) were generated with timetree (80). We used codeml models M0 to estimate one $D_N/D_S$ for each gene and codeml models M1, M2, M7 and M8 for likelihood ratio tests (LRTs) and estimation of post. mean $D_N/D_S$. Site models allow the $D_N/D_S$ ratio to vary for sites across the gene. Models 1 (nearly neutral) and 2 (positive selection), 7 (beta) and 8 (beta&ω) are nested models designed to test for signatures of positive selection. Since only models 2 and 8 allow for sites under positive selection, we can use LRTs and the LRT statistic (twice the log-likelihood difference between the two compared models ($2\Delta\ell$)), and a chi-square test to see which model is favored. LRTs to detect signatures of positive selection were performed using codeml.

**Structure-informed PAML.** For structure-informed analyses we used the approximate post. mean $D_N/D_S$ ratio (ω) computed using the Bayes empirical Bayes (BEB) method under the codeml model M8 as an estimate of selection. We mapped these estimates as attributes onto the Alphafold protein structures using ChimeraX (67) and custom R scripts. All Alphafold protein models downloaded from the database were version 4. A similar approach has previously been implemented in the Selecton server (19). To make visual inspection more straightforward we use a fixed color scheme across all structures with mean $D_N/D_S$ capped at 1.5. Since post. mean $D_N/D_S$ estimates for single residues are based on few datapoints, their accuracy is limited and highly sensitive to the number of sequences, distance between sequences, and selection. To see how robust estimation is within our dataset, we compared post. mean $D_N/D_S$ estimates generated under M8 using the empirical Bayes method with estimates generated under different codeml models (M1, M2, M7 and M8 using naïve empirical Bayes or BEB) (***SI Appendix* Fig. S1*F***). We used the post. mean $D_N/D_S$ estimates generated under M8 for further analyses. For consistency with previous publications, residues with a post. probability higher than 0.9 that $D_N/D_S$ is greater than 1 were considered significant. While empirical Bayesian methods suffer from inaccuracy in small datasets, we deem it useful to distinguish between strong purifying selection and loss of purifying selection, especially when combined with structural and functional information about the amino acid residues to judge how much purifying selection to expect based on the mechanistic/biochemical context. All custom R scripts required for the analyses are available on Figshare.

**PAML fixed-site models.** To perform statistical tests comparing selection rates between paralogous proteins or between specified domains or sections of proteins, we used partition gene models using the PAML fixed-site models as previously described (68). Protein domain boundaries were obtained from Interpro (69).

**Analyses of mouse population genomic data.** Population genomic data for 23 mouse strains were downloaded as VCF files from (28). The population samples were from France (14, 15B, 16B, 18B), Iran (413, 416, AH15, AH23, JR11, JR15), Kazakhstan (AL16, AL19, AL1), Czech Republic (CR12, CR13, CR16, CR23, CR25, CR29), India (H12, H36), and Germany (TP121B, TP1). The variants were extracted from the VCF file for each strain and reconstituted to the mm10 reference genome using the *vcf_to_fasta.sh* script, which utilizes the GATK *FastaAlternateReferenceMaker* tool (81). The mm10 RefSeq genes were then extracted from the variant reconstituted genomes using the bedtools *getfasta* tool (82) Next, we performed McDonald-Kreitman tests (83) with the *MK.pl* script, using *M. spretus* (SP36 strain) as an outgroup. The direction of selection metric (84) was then calculated using the following formula:

$$DoS = \frac{D_N}{D_N + D_S} - \frac{P_N}{P_N + P_S}$$

where $D_N$ and $D_S$ are divergent non-synonymous and synonymous sites, respectively, and $P_N$ and $P_S$ are polymorphic non-synonymous and synonymous sites. We used DNAsp to estimate direction of selection (29). All custom scripts required for the analyses are available on Figshare.

**Protein structure prediction and visualization.** Protein structural models were downloaded from the Alphafold database (21). Models were not available for *T. guttata* RAD50, DMC1, HFM1, BLM, RNF212B, MLH3 and MUS81, these proteins were modelled using the ColabFold workbooks and the UniProt amino acid sequence (70). In this case the first model was chosen for each protein. Protein structure visualization and analyses of solvent exposed surface areas (SESA) were performed in UCSF Chimera (71) or ChimeraX (67). We calculated a relative SESA using theoretical maximum values for each amino acid, as previously described (72). For the yeast Spo11 core complex we used a dimer prediction based on the published cryo-EM structure (38). For other protein complexes, we used predictions generated by the Alphafold3 online server (42). Interaction surfaces were identified using ChimeraX by selecting residues that bury ≥ 15 $Å^2$ of solvent-accessible surface in the interface between chains. We then grouped all residues into three categories: buried (relSESA < 0.3), surface (relSESA ≥ 0.3 in the monomeric form), and contact surface (relSESA ≥ 0.3 in the monomeric form, with interactions in the complex structure). Across the four lineages and four complexes, we classified a total of 3674 contact surface residues. We used the ChimeraX *swapaa* function to analyze how the different amino acid combinations found in different rodent species affect these interactions. For pairwise structure alignments we used the FATCAT server with default parameters using the flexible alignment model (73). Only high-confidence alignments were included for further analyses (Tm-score>0.5 and sufficient coverage of the whole protein sequence). Our final dataset included comparisons between RAD51 and DMC1 as well as MSH2 and MSH4. Calculation of the difference in post. mean $D_N/D_S$ between structurally matched residues was performed in R using custom scripts. Protein domain annotations throughout the manuscript are from Interpro (69). All scripts and resources are available from Figshare.

**Cloning and strain construction.** All strains were SK1 derivatives, as detailed in *SI Appendix* Table S1. The following alleles have been published previously: spore-autonomous fluorescent markers *THR1::m-Cerulean-TRP1*, *CEN8::tdTomato-LEU2*, and *ARG4::GFP*-URA3* (48). To replace the endogenous copies of *MSH4 and MSH5*, *MSH4* or MSH5 sequences were cloned into a plasmid containing a FLAG tag and HpHMX resistance cassette by PCR. Subsequent gene replacements were then performed using In-Fusion cloning (Takara). Linearized vectors were then transformed into the wild-type SK1 background using standard cloning.

**Protein analyses by immunoblot.** Samples for protein analyses were prepared from synchronously sporulating yeast cultures. In brief, diploid colonies were grown on YP-glycerol plates (2% peptone, 1% yeast extract, 2% glycerol, 2% agar) for 48 hrs at 30°C. Cells were then transferred onto YPD plates (2% peptone, 1% yeast extract, 2% dextrose, 2% agar) and grown to form a small patch (overnight, 30°C). Cells were then transferred to YPD plates and grown into a lawn (overnight, 30°C), which was then used to inoculate pre-sporulation medium YP2%KAc (2% peptone, 1% yeast extract, 2% potassium acetate) to $OD_{600}$ ~0.3. Cells were grown for 11 hr at 30°C, washed with sporulation medium (SPM, 2% potassium acetate) and inoculated into SPM to $OD_{600}$ ~3.5. This time point was defined as t = 0 in all meiotic experiments. Ten ml of culture were taken to prepare TCA protein extracts. Samples were disrupted using a bead beater and glass beads in 10% trichloroacetic acid. Precipitates were resuspended in 2× NuPAGE sample buffer and neutralized with 1 M Tris base. Samples were heated (95°C) and cleared by centrifugation. Samples were loaded on NuPAGE 4–12% Bis-Tris (Invitrogen). Gels were blotted on PVDF membranes using semi-dry transfer (GE Healthcare). HRP-conjugated anti-FLAG antibody (Sigma A8592) was used for detection of Msh4, HRP-conjucated anti-myc antibody (9E10, Abcam ab32) was used for the detection of Msh5. Rat anti Tubulin Alpha (Biorad, MCA78G) antibody was used for loading controls. Blots were imaged on a BioRad ChemiDoc system.

**Spore viability.** To assess spore viability, diploid yeast strains were sporulated on SPM plates (2% potassium acetate) overnight at 30°C. Tetrads were treated with Zymolyase 100T (Zymo research) for 10 min at room temperature to disrupt the cell wall. Spore viability was determined by microdissection using a Singer SporePlay+ Dissection instrument. Spore viability was quantified as the number of viable spores per dissected tetrad (0–4).

We assessed genotype effects using a beta-binomial generalized linear model. An ordinary binomial model would assume a single, fixed probability of spore death per genotype, with each spore an independent trial of that probability. We considered this assumption to be inappropriate for tetrad data: the four spores within a tetrad share a single meiotic division, so any factor that affects that division (e.g., a nondisjunction event, or, in this experiment, how well a given heterologous protein functions in that particular cell) affects the fates of all four spores jointly rather than each spore separately. The result is overdispersion: variance in viability across tetrads exceeds what a binomial model, applied at the level of individual spores, would predict. The beta-binomial model addresses this by adding a free parameter that governs how much the spore-death probability is allowed to vary from tetrad to tetrad, rather than treating that probability as fixed within a genotype.

The beta-binomial model was fit to the observed pattern of tetrad counts (how many tetrads had 0 viable spores, how many had 1, and so on, for each genotype) to determine a death-probability estimate per genotype and its associated uncertainty, plus a shared dispersion parameter. Models were fit by maximum likelihood using the glmmTMB package (version 1.1.10) in R (version 4.4.1), which finds these parameter values numerically via nonlinear optimization. Genotype comparisons were then made by comparing these fitted probabilities, and by using an LRT to test whether a model that allows genotypes to differ explains the data significantly better than one that does not.

Pre-specified contrasts, based on expected evolutionary ordering of heterologous complementation defects, were evaluated using the following hierarchical gatekeeping strategy. We first tested whether any heterologous *MSH4* allele differed from the reintegration control (omnibus test). If significant ($\alpha = 0.05$), we tested each heterolog individually against the reintegration control using one-sided tests with Holm-Bonferroni adjustment. When two or more heterologs showed significant defects, we performed stepwise contrasts (*S. arboricola* vs. *S. paradoxus*; *S. bayanus* vs. *S. arboricola*) to assess evolutionary ordering. Control strain comparisons (wild type, *msh4*Δ, reintegration) were evaluated separately to validate experimental methods. Confidence intervals for observed viability were estimated by bootstrap resampling of tetrads (5,000 replicates). We also assessed MI-NDJ signatures by comparing the frequency of tetrads with 0 or 2 viable spores across genotypes using logistic regression with the same hierarchical testing structure.

The *MSH5* rescue experiments were treated as a separate hypothesis family from the primary *MSH4* heterology comparisons, with multiple-testing correction applied independently within that family. Three comparisons were made in which the endogenous *MSH5* was replaced with *MSH5* from *S. cerevisiae*, *S. arboricola*, or *S. bayanus* in a strain that also had the cognate MSH4 integrated from the same species. That is, *S. bayanus MSH5* was introduced into a strain that already had *S. bayanus MSH4*, and so on. We used two-sided tests (with Holm-Bonferroni across the three tests; **Datasets S7–9**) because there was no a priori prediction as to whether introducing the cognate *MSH5* would improve or worsen meiotic processes, as explained in the main text.

Raw data and custom R scripts for the statistical analysis are available at Figshare.

**Analysis of recombination.** The spore-autonomous fluorescence recombination analysis was performed as described previously (48). Yeast cells were synchronized as described above and inoculated into SPM to $OD_{600}$ ~3.5. Cells were incubated at 30°C for ~48 hrs and subsequently imaged on a Zeiss Axio Observer Z1 Marianas Workstation, equipped with an ORCA-Flash 4.0 camera, illuminated by an X-Cite 120 PC-Q light source, with 60× air objective. Marianas Slidebook (Intelligent Imaging Innovations) software was used for acquisition. The pattern of fluorescence in the tetrads was manually scored using Fiji. Only tetrads with four spores and each fluorescence marker occurring in two spores were included in the final analysis. Genetic distances (in centiMorgans, cM) were calculated using the Perkins equation (85):

$$cM = \frac{100 * (6 * NPD) + TT}{2 * (PD + TT + NPD)}$$

This equation corrects for multiple crossovers and provides an unbiased estimate of genetic distance from tetrad data. MI-NDJ events were excluded from cM calculations and analyzed separately. We compared genotypes using Fisher's exact tests, which make no distributional assumptions and are robust to overdispersion. Planned contrasts followed the same hierarchical structure used for viability analysis and were applied independently to each interval and to MI-NDJ. Confidence intervals for cM were obtained by bootstrap resampling of tetrad counts (10,000 replicates). Additionally, complementary to the Fisher tests, directional differences (ΔcM) were assessed using one-sided bootstrap tests reflecting the *a priori* expectation that complementation defects increase with evolutionary distance. The fluorescent reporter data for the *MSH5* complementation experiment were analyzed using two-sided tests with Holm-Bonferroni correction for the reasons described above for the tetrad viability data.

### Supplemental Figures

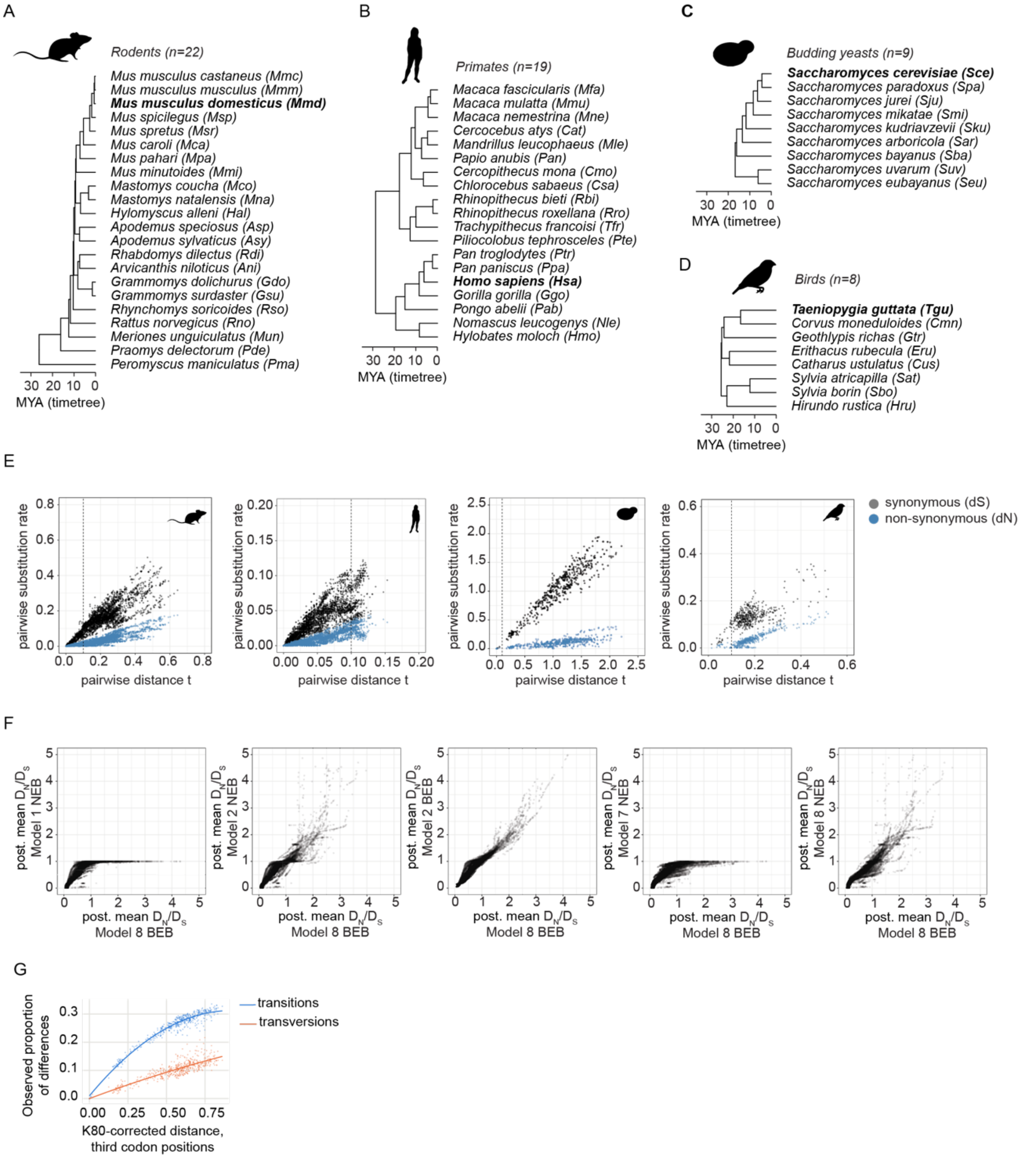


**Fig. S1.** Species trees and pairwise substitution rates. (*A*-*D*) Species trees with bold text indicating representative species for which Alphafold models were obtained. Trees were generated using timetree (79). MYA, million years ago. (*E*) Pairwise synonymous (grey) and non-synonymous (blue) substitution rates plotted against the pairwise distance t for all genes. The vertical dashed lines in each plot indicate t = 0.1. (*F*) Correlation of post. mean $D_N/D_S$ estimates generated for each codon under different codeml models. Note that Models 1 and 7 do not allow for sites with $D_N/D_S > 1$. (*G*) Analysis of saturation at synonymous sites in the yeast dataset (see ***SI Appendix Materials and Methods***)

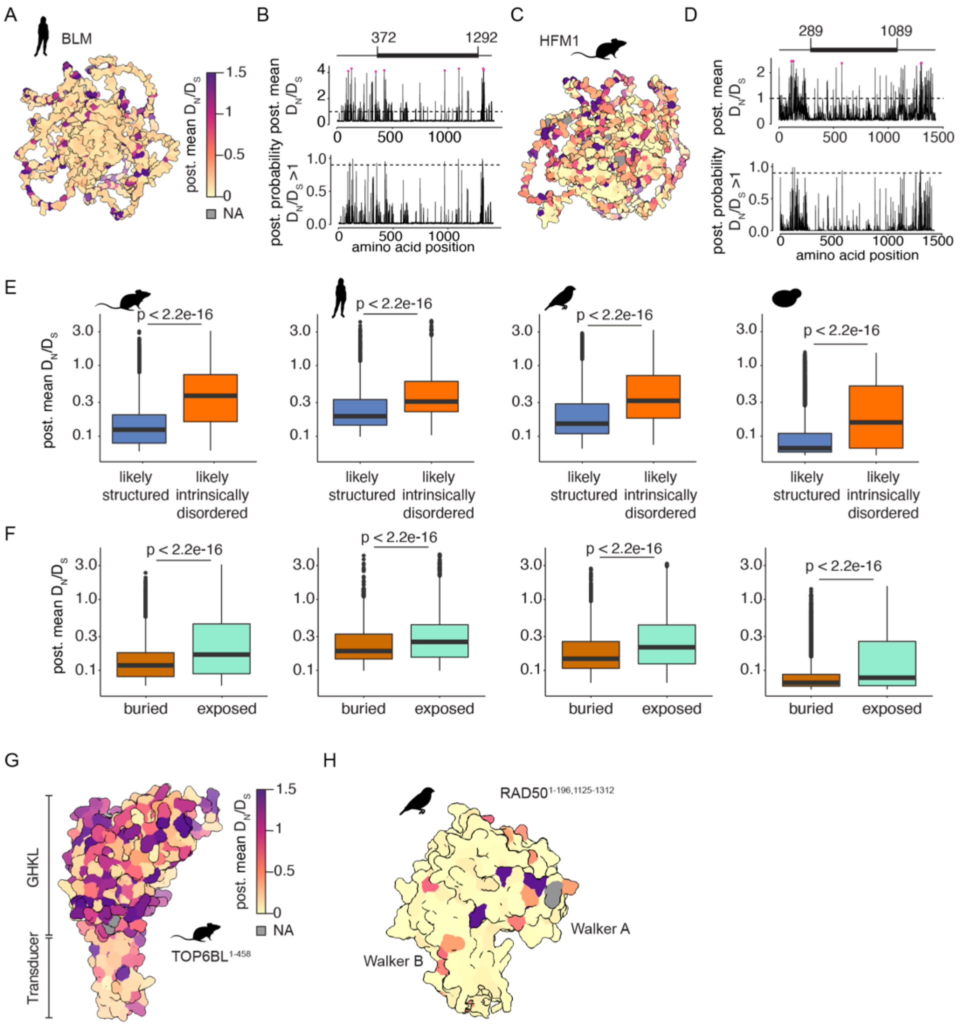

**Fig. S2.** Structural constraints influence selection rates. (*A*-*D*) Analysis of primate BLM (*A*,*B*) and rodent HFM1 (*C*,*D*). Structural models of human (*A*) or mouse (*C*) proteins are colored according to post. mean $D_N/D_S$. The graphs in panel *B* and *D* show post. mean $D_N/D_S$ (top) and post. probability for positive selection (bottom) for each residue. Residues with a post. probability > 0.9 are shown in pink. Horizontal dashed lines indicate $D_N/D_S = 1$ or post. probability = 0.9. The schematics above in panel *B* and *D* show positions of the central structured domains (thick lines) and unstructured N- and C-terminal domains (thin lines). (*E*,*F*) For each lineage, post. mean $D_N/D_S$ values for residues categorized according to pLDDT as likely structured or likely intrinsically disordered (*E*) or categorized according to rel. SESA as buried or surface (*F*). (*G*,*H*) Post. mean $D_N/D_S$ mapped onto AlphaFold models of mouse TOP6BL (*G*) and the zebra finch RAD50 head domain (*H*).

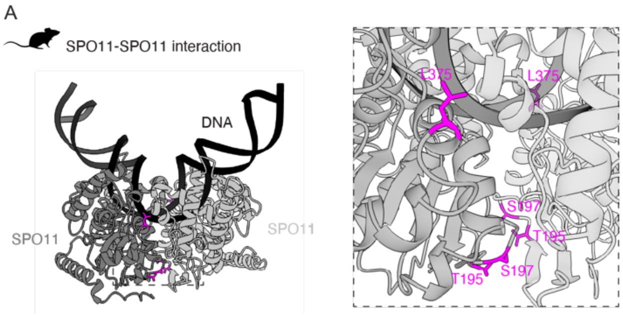


**Fig. S3.** Variation of the SPO11–SPO11 dimer interface in rodents. (*A*) Overview and detailed view of the Alphafold structure of two SPO11–TOP6BL complexes bound to bent DNA. TOP6BL is omitted to highlight the SPO11 dimer configuration. Residues at the dimer interface that showed post. mean $D_N/D_S > 1$ (T195, S197 and L375) are highlighted in magenta.

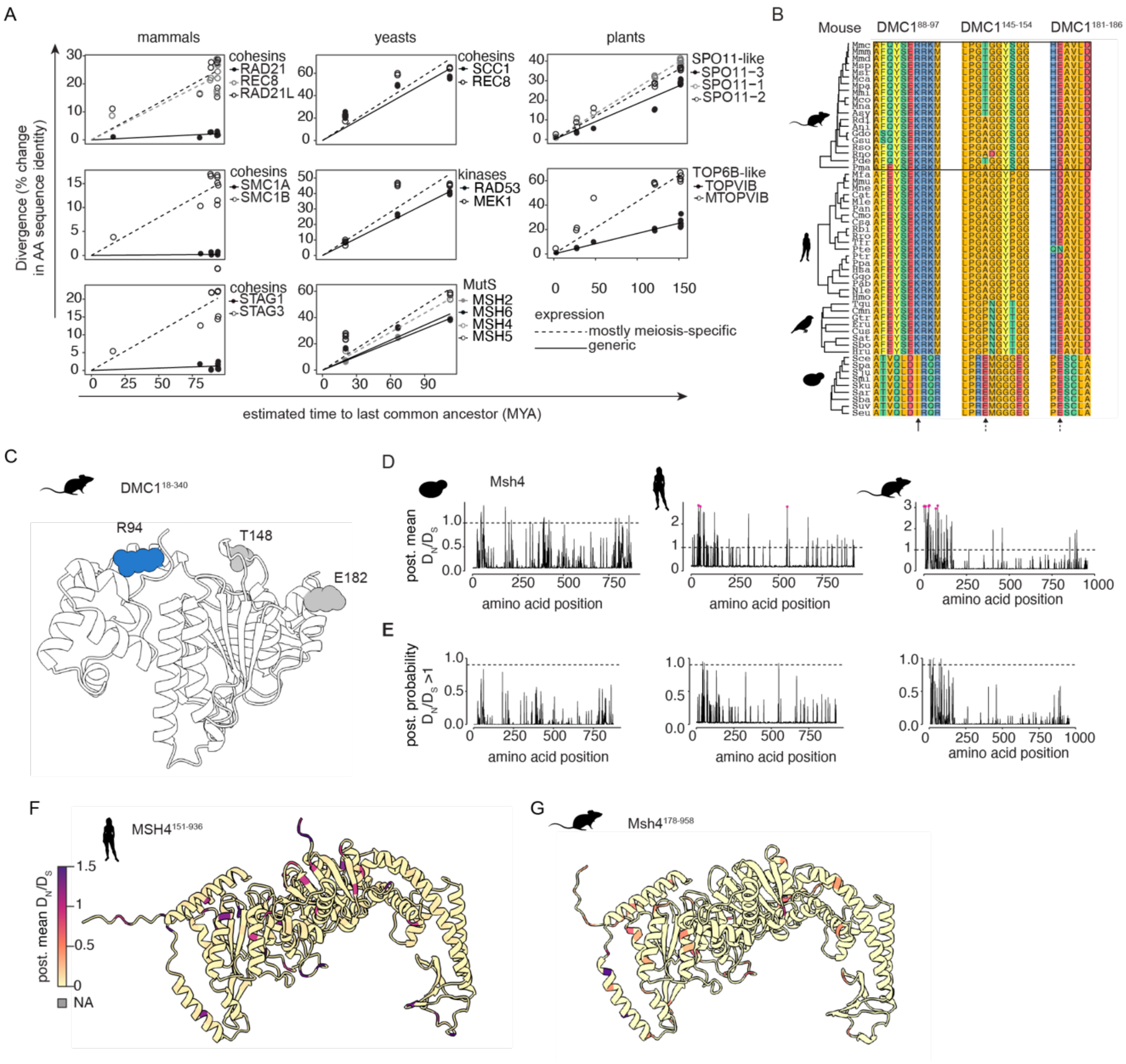

**Fig. S4.** Paralog comparisons. (*A*) Pairwise divergence for additional selected paralogous protein pairs presented as described for **Fig. 4*A***. (*B*) Partial alignment of DMC1. Black rectangle highlights the rodent lineage in which we initially observed relaxed purifying selection. Solid arrows indicate residues with post. mean $D_N/D_S$ >1 and dashed arrows show residues with $D_N/D_S$ just below 1. (*C*) Mouse DMC1. The blue residue (R94) shows post. mean $D_N/D_S$ > 1; gray residues have $D_N/D_S$ just below 1 (T148, E182). (*D*,*E*) Post. mean $D_N/D_S$ (*D*) and post. probability for positive selection (*E*) for each residue in MSH4 in three lineages. Residues with a post. probability > 0.9 are shown in pink. Horizontal dashed lines indicate $D_N/D_S$ = 1 or post. probability = 0.9. (*F*,*G*) Post. mean $D_N/D_S$ mapped onto the AlphaFold models of *H. sapiens* MSH4 (*F*) and *M. musculus* MSH4 (*G*).

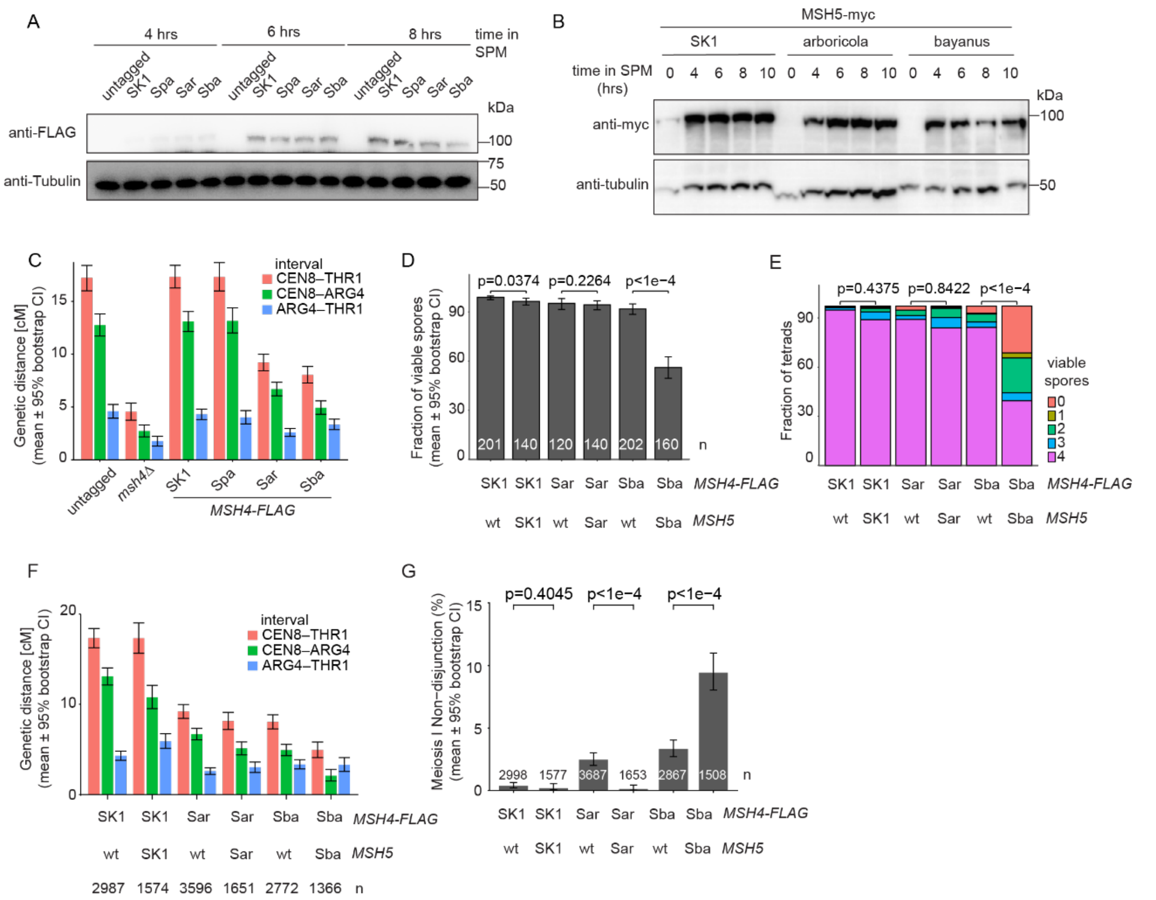


**Fig. S5.** Functional relevance of Msh4 and Msh5 sequence variation in budding yeasts. (*A*) Immunoblot analysis of Msh4-FLAG expression at the indicated times during meiosis. (*B*) Immunoblot analysis of Msh5-myc expression at the indicated times during meiosis. Endogenous *MSH5* in *S. cerevisiae* was replaced with myc-tagged versions of *MSH5* from *S. cerevisiae* (SK1 strain), *S. arboricola*, or *S. bayanus*. (*C*) Genetic distance data for three intervals (*CEN8–THR1* and its two component intervals) related to **Fig. 5*E***. Bars indicate mean genetic distances and error bars represent 95% bootstrap confidence intervals. At least two independent experiments were performed per genotype. (D–G) Effects of combining heterologous Msh4 with cognate Msh5 heterologs from the same species. (*D*) Spore viability. Data for the heterologous *MSH4* strains that contain the endogenous *MSH5* are re-used from **Fig. 5*B***. Bars show the means and error bars indicate 95% bootstrap confidence intervals. The total number of tetrads scored is indicated for each genotype. Two-sided p-values are from beta-binomial generalized linear model testing after Holm-Bonferroni correction. (*E*) The number of viable spores per tetrad was scored to detect signatures of MI-NDJ events. Stacked bar plots show the mean number of viable spores per tetrad for the same tetrad data as in panel *D*. P-values as in panel *D*. (*F*) Genetic distance data calculated using the Perkins equation for the indicated genotypes. Bars indicate mean genetic distances and error bars represent 95% bootstrap confidence intervals. The total number of tetrads for each genotype is indicated excluding MI-NDJ events. At least two independent experiments were performed per genotype. Fisher's test two-sided p-values after Holm-Bonferroni correction are shown. (*G*) MI-NDJ was measured from the same tetrad data as in panel *F*. Bars show the mean, error bars indicate 95% Wilson binomial confidence intervals.

Fisher's test two-sided p-values after Holm-Bonferroni correction are shown. *Note:* Overall spore viability was slightly but significantly decreased in the *S. cerevisiae MSH4* + *S. cerevisiae MSH5* reintegration strain (panel *D*), but we consider this unlikely to be biologically meaningful given that significant differences were not seen in analysis of the tetrad viability MI-NDJ signature (*E*) or fluorescence-based crossover or MI-NDJ measurements (*F,G*, **Dataset S8**). MI-NDJ in the fluorescence assay appeared to improve with the *S. arboricola* Msh5 (*G*), but this was not supported by spore viability (which reports on whole-genome NDJ, panel *E*) or crossing over (*F*, **Dataset S8**). Our overall conclusion is that adding in *S. arboricola* Msh5 does not substantially alter *S. arboricola* Msh4 behaviour in the cross-complementation test, but we cannot exclude quantitatively small effects.

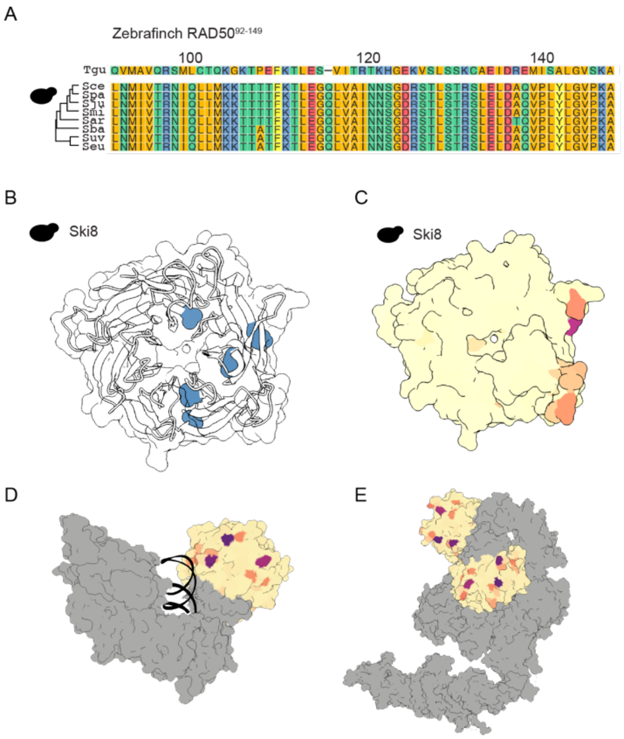


**Fig. S6.** Variation within Ski8 in budding yeasts. (*A*) Partial sequence alignment of Ski8 in budding yeasts and Zebrafinch for comparison. (*B*) Variation of structured residues. A transparent surface and ribbon representation of budding yeast Ski8 is shown. The five blue residues (H58, L81, V142, G198, V382) show a subtle relaxation from purifying selection (post. mean $D_N/D_S$ > 0.4) despite strong structural constraints. (*C*) Variation of surface residues. Post. mean $D_N/D_S$ is mapped onto a surface representation of the AlphaFold model of budding yeast Ski8. (*D*,*E*) Orientation of Ski8 in the context two complexes: the Spo11 core complex (monomeric cryo-EM structure from (38)) in (*D*) and the Ski2–Ski3–Ski8 complex (PDB: 4BUJ) in (*E*). Only Ski8 is colored by post. mean $D_N/D_S$ to highlight the position of the diverging surface.

**Table S1. List of yeast strains.**

| Strain | MAT | Genotype | Source/Ref. |
|---|---|---|---|
| SKY7454, SKY7604 | *a/α* | wild type SK1 diploid | This study |
| SKY7421 | *a/α* | *MSH4(SK1)-FLAG-HpHMX4* | This study |
| SKY7422 | *a/α* | *MSH4(paradoxus)-FLAG-HpHMX4* | This study |
| SKY7424 | *a/α* | *MSH4(arboricola)-FLAG-HpHMX4* | This study |
| SKY7425 | *a/α* | *MSH4(bayanus)-FLAG-HpHMX4* | This study |
| SKY7481 | *a/α* | *msh4Δ::HpHMX4* | This study |
| SKY7445 | *a/α* | *THR1/THR1::m-Cerulean-TRP1 CEN8/CEN8::tdTomato-LEU2 ARG4/ARG4::GFP*-URA3* | This study |
| SKY7446 | *a/α* | *MSH4(SK1)-FLAG-HpHMX4 THR1/THR1::m-Cerulean-TRP1 CEN8/CEN8::tdTomato-LEU2 ARG4/ARG4::GFP*-URA3* | This study |
| SKY7447 | *a/α* | *MSH4(paradoxus)-FLAG-HpHMX4 THR1/THR1::m-Cerulean-TRP1 CEN8/CEN8::tdTomato-LEU2 ARG4/ARG4::GFP*-URA3* | This study |
| SKY7450 | *a/α* | *MSH4(arboricola)-FLAG-HpHMX4 THR1/THR1::m-Cerulean-TRP1 CEN8/CEN8::tdTomato-LEU2 ARG4/ARG4::GFP*-URA3* | This study |
| SKY7451 | *a/α* | *MSH4(bayanus)-FLAG-HpHMX4 THR1/THR1::m-Cerulean-TRP1 CEN8/CEN8::tdTomato-LEU2 ARG4/ARG4::GFP*-URA3* | This study |
| SKY7452 | *a/α* | *msh4Δ::HpHMX4 THR1/THR1::m-Cerulean-TRP1 CEN8/CEN8::tdTomato-LEU2 ARG4/ARG4::GFP*-URA3* | This study |
| SKY7572 | *a/α* | *MSH4(SK1)-FLAG-HpHMX4, MSH5(SK1)–KanMX* | This study |
| SKY7573 | *a/α* | *MSH4(SK1)-FLAG-HpHMX4, MSH5(SK1)–KanMX, THR1/THR1::m-Cerulean-TRP1 CEN8/CEN8::tdTomato-LEU2 ARG4/ARG4::GFP*-URA3* | This study |
| SKY7580 | *a/α* | *MSH4(arboricola)-FLAG-HpHMX4, MSH5(arboricola)–KanMX* | This study |
| SKY7581 | *a/α* | *MSH4(arboricola)-FLAG-HpHMX4, MSH5(arboricola)–KanMX, THR1/THR1::m-Cerulean-TRP1 CEN8/CEN8::tdTomato-LEU2 ARG4/ARG4::GFP*-URA3* | This study |
| SKY7588 | *a/α* | *MSH4(SK1)-FLAG-HpHMX4, MSH5(SK1)–KanMX* | This study |
| SKY7589 | *a/α* | *MSH4(arboricola)-FLAG-HpHMX4, MSH5(arboricola)–KanMX, THR1/THR1::m-Cerulean-TRP1 CEN8/CEN8::tdTomato-LEU2 ARG4/ARG4::GFP*-URA3* | This study |
| SKY7594 | *a/α* | *MSH4(SK1)-FLAG-HpHMX4, MSH5(SK1)–myc10–HpHMX4* | This study |
| SKY7599 | *a/α* | *MSH4(arboricola)-FLAG-HpHMX4, MSH5(arboricola)–myc10–HpHMX4* | This study |

SK1 background *ho::LYS2, lys2, ura3, leu2, trp1::hisG*
